# Cellular and Immune Adaptations at the Maternal–Fetal Interface in Bats

**DOI:** 10.1101/2025.05.15.654282

**Authors:** Allyson Caldwell, Liheng Yang, Rebecca L. Casazza, Rizban E. Worota, Cole McCutcheon, Patrick S. Creisher, Erika Zhan, Clara Reasoner, Ashley Higgins, Tony Schountz, Carolyn B. Coyne

## Abstract

Bats maintain pregnancy despite extended gestation relative to other small mammals, high fetal investment, recurrent pathogen exposure, and the metabolic demands of flight. These physiological extremes likely drive unique adaptations in placental function and maternal-fetal immune regulation, yet the cellular and molecular basis of these adaptations remains largely unknown. Here, we mapped the cellular landscape of the Jamaican fruit bat (*Artibeus jamaicensis*) placenta using single-nucleus RNA sequencing (snRNA-seq), integrated with histological and immunohistochemical analyses. We identified diverse trophoblast, stromal, and immune populations at the maternal-fetal interface, including specialized macrophages expressing pregnancy-associated signaling molecules. Trajectory analysis revealed dynamic trophoblast differentiation through proliferative, invasive, and syncytial states. To model these processes *in vitro*, we derived trophoblast and decidual gland organoids from matched tissues, which recapitulated key *in vivo* cell types and lineage trajectories. Cross-species transcriptomic comparisons with human and mouse placentas uncovered bat-specific gene programs in trophoblasts, fibroblasts, and immune cells. Notably, bat trophoblast organoids exhibited attenuated antiviral signaling compared to their human counterparts, suggesting species-specific modulation of innate immunity at the maternal-fetal interface. These findings define cellular strategies that support pregnancy under extreme physiological conditions and establish a framework for investigating the evolution of placental adaptations across mammals.

Graphical abstract:
Using single-nucleus RNA sequencing (snRNA-seq) and placental organoids, we reveal unique trophoblast differentiation, immune cell specialization, and expanded stromal populations in Jamaican fruit bat placentas. Cross-species comparisons underscore bat-specific evolutionary adaptations critical for sustaining pregnancy amid extreme metabolic and immunological challenges.

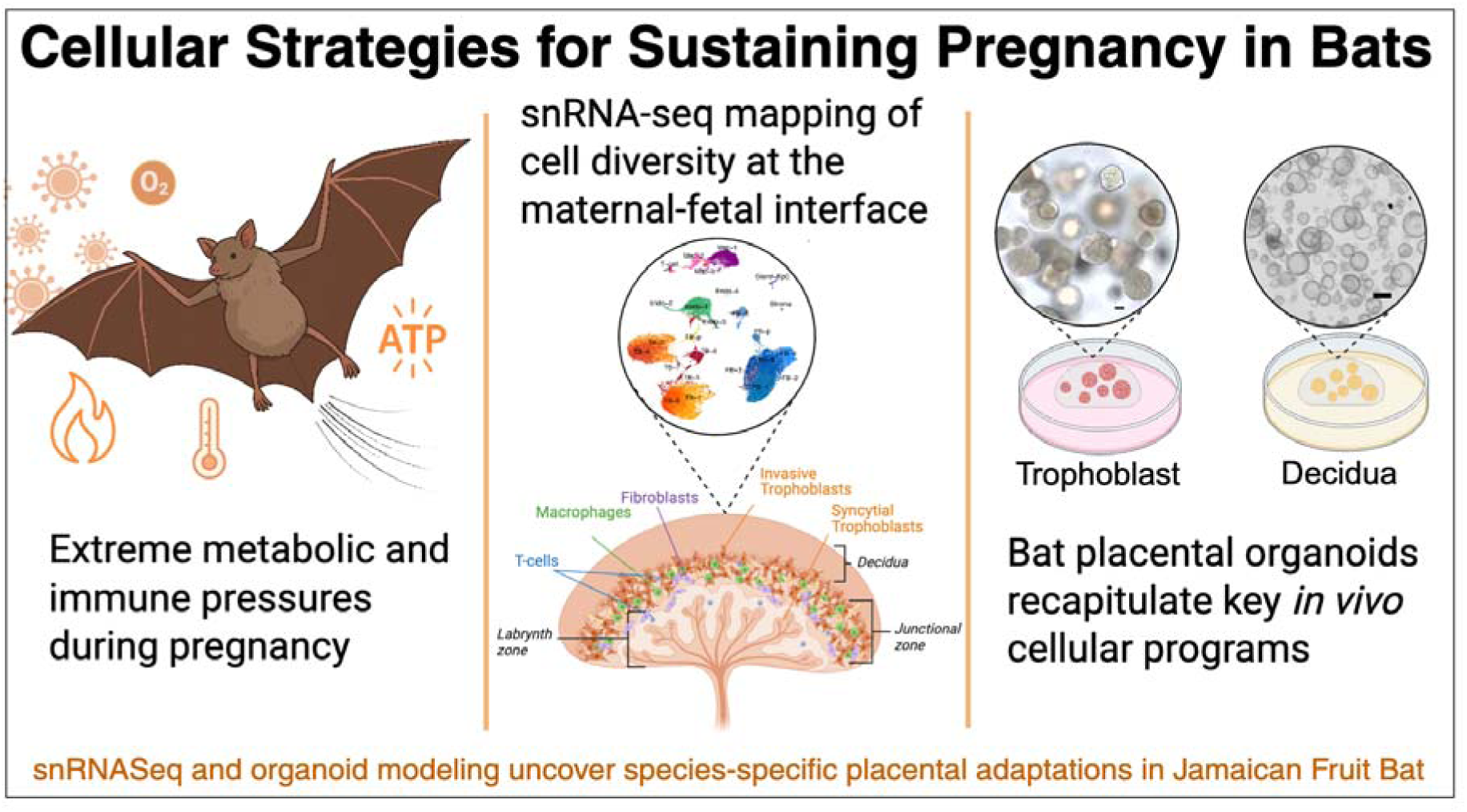

**Summary:** Pregnancy in bats occurs under a combination of physiological stressors rarely encountered in other mammals, including extended gestation relative to other small mammals, elevated metabolic load, temperature fluctuations associated with flight, and constant microbial exposure. To investigate how the placenta adapts to these challenges, we profiled Jamaican fruit bat (*Artibeus jamaicensis*) placenta using single-nucleus RNA sequencing and tissue-derived organoid models. This analysis revealed diverse trophoblast, stromal, and immune cell populations with bat-specific transcriptional programs. We identified dynamic trophoblast differentiation trajectories, fibroblasts with hybrid adventitial-neuronal signatures, and macrophages expressing pregnancy-associated molecules typically restricted to trophoblasts. Cross-species comparisons with human and mouse placentas uncovered both conserved cell states and lineage-specific adaptations, providing insight into the evolution of placental function and immune regulation across mammals.

**Highlights:**

- Single-nucleus RNA-seq defines cell types and differentiation pathways in the bat placenta
- Bat trophoblast and decidua gland organoids recapitulate *in vivo* lineage trajectories and cell signatures
- Bat-specific transcriptional programs revealed across trophoblast, stromal, and immune lineages
- Bat trophoblasts exhibit attenuated antiviral responses compared to human counterparts

## Introduction

Mammalian pregnancy requires extensive physiological, cellular, and immune adaptations to support fetal development while preserving maternal health. These adaptations vary widely across species, shaped by factors such as gestation duration, litter size, maternal immune architecture, and ecological pressures. Central to a successful pregnancy is the placenta, a transient organ that mediates nutrient and gas exchange, hormone production, and immunological tolerance at the maternal-fetal interface. While all placentas serve these core functions, they exhibit striking diversity in structure, cellular composition, and maternal-fetal exchange strategies, reflecting millions of years of evolutionary innovation across divergent mammalian lineages^1,2^.

Among mammals, bats present a compelling system for investigating how the placenta adapts to physiological extremes. As the second-largest mammalian order with over 1,400 species, bats exhibit remarkable variation in placental morphology and reproductive strategies^3^. Many species sustain prolonged gestation relative to body size, with some employing delayed implantation or developmental suspension to synchronize birth with environmental cues. In contrast to most other small mammals that produce short-gestation litters, many bats give birth to a single, highly developed offspring after extended intrauterine development. These strategies suggest that bat placentas must support sustained fetal growth under unique mechanical and metabolic constraints.

In parallel, bats face physiological and immunological challenges that place unique demands on placental function. Sustained flight imposes high metabolic rates and oxidative stress, whereas torpor introduces fluctuating thermal and hemodynamic states^4^. Despite dense roosting and high population mobility that elevate pathogen exposure, bats serve as natural reservoirs for a wide range of zoonotic viruses including coronaviruses, filoviruses, and henipaviruses, yet show no signs of disease^5–8^. Multiple studies have revealed that bats exhibit dampened activation of key immune signaling pathways, including interferon responses, NF-κB signaling, and inflammasome activation, despite maintaining constitutive expression of select antiviral genes^9–11^. This tolerance to persistent viral presence raises critical questions about how similar regulatory strategies may function at the maternal-fetal interface, where immune equilibrium is essential for successful gestation. How these systemic immune adaptations are integrated into placental biology remains unknown.

Previous studies on bat placentation have primarily focused on morphological descriptions. Early work characterized the fetal membranes and vascular structures in the vampire bat (*Desmodus rotundus*), revealing invasion patterns distinct from those of primates and rodents^12^. Subsequent studies reported placental variation across species, including both endotheliochorial and hemochorial types^13–15^. Most microbats possess hemomonochorial placentas, with a single syncytial trophoblast layer interfacing directly with maternal blood, supported by underlying cytotrophoblasts. Some species exhibit dynamic changes in placental structure throughout gestation, transitioning from endotheliochorial to hemodichorial forms^16^. However, the cellular and molecular mechanisms governing bat placental development, such as trophoblast lineage specification, immune cell recruitment, and nutrient transport regulation, remain largely unexplored.

Trophoblasts are the defining epithelial lineage of the placenta, but their differentiation trajectories and functions vary widely across mammals. In hemochorial placentas, such as those in humans, cytotrophoblast progenitors give rise to the multinucleated syncytiotrophoblast that mediates nutrient, gas, and waste exchange, as well as to invasive extravillous trophoblasts that remodel the maternal vasculature. Additional placental cell types including stromal fibroblasts, endothelial cells, glandular epithelium, and diverse immune populations act together to support implantation, fetal tolerance, and tissue remodeling. Although these lineages have been well defined in human and murine placentas, their composition, developmental dynamics, and transcriptional programs remain uncharacterized in bats. As a result, the cellular basis for placental specialization in this lineage, including how bats sustain gestation, tolerate fetal antigens, and resist infection at the maternal-fetal interface, remains largely unknown.

To address these knowledge gaps, we combined single-nucleus RNA sequencing (snRNA-seq) and tissue stem cell-derived organoid models to map the cellular and molecular architecture of the Jamaican fruit bat (*Artibeus jamaicensis*) placenta. High-resolution snRNA-seq enabled us to define the full spectrum of trophoblast, stromal, endothelial, and immune populations at the maternal-fetal interface, while matched organoid models captured key trophoblast lineages and allowed *in vitro* reconstruction of trophoblast differentiation programs. This integrative approach revealed bat-specific cellular adaptations that support prolonged gestation and modulate maternal immune responses. Cross-species comparisons with human and mouse placentas uncovered both conserved cell states and divergent transcriptional programs, providing new insights into the evolution of placental biology, trophoblast specialization, and maternal-fetal immune strategies across mammals.

## Results

### Isolation and characterization of Jamaican fruit bat placenta

Placentas were collected from five visibly pregnant adult female Jamaican fruit bats (Jfb) that were euthanized in accordance with approved animal protocols. Immediately following euthanasia, the reproductive tracts were dissected and the placentas isolated for downstream analyses. Gestational stage was estimated based on previously established morphological criteria for related phyllostomid species^17^. These included relative fetal crown–rump length (CRL), degree of wing digit separation, wing membrane translucency, and the ratio of placental disc size to fetal size, as described in prior studies of chiropteran development. In particular, reopening of the eyes, increased pigmentation in skin and claws, elongation of the digits, and presence of a prominent, ribbed nose-leaf indicated that all fetuses were in stage 21-24 of development, with one placenta determined to be in the fetal period of development (Plac1, Stage 24) based on fetal CRL, skin wrinkling, and pigmentation (**Figure S1A, S1B)**.

To characterize the morphology of the Jfb placenta, we fixed placental tissue for histological analysis and processed it alongside late-gestation (∼E17.5) mouse placentas for comparison. Hematoxylin and eosin staining revealed a large, highly vascularized labyrinth that occupied approximately two-thirds of the bat placental section. This structure was morphologically analogous to the murine labyrinth zone, suggesting a conserved organization of the exchange interface between species (**Figure 1A-1F**). Immunohistochemistry (IHC) using either a pan-cytokeratin (Pan-CK) antibody or a cytokeratin 18 (KRT18)–specific antibody was performed to identify epithelial and trophoblast cell populations. Cytokeratin staining revealed two distinct trophoblast regions: a large, vascularized labyrinth zone, and a narrow band of morphologically distinct trophoblasts situated between the labyrinth and the decidua. We refer to this intermediate region as the "junctional zone," consistent with the nomenclature used to describe similarly positioned trophoblasts in the mouse placenta (**Figure 1A-1F**, middle panels and **Figure S1E**). The mouse junctional zone is comprised of unique trophoblast types including spongiotrophoblasts, glycogen trophoblasts, and trophoblast giant cells (TGCs)^18^. Although similar in location, the trophoblasts in the bat junctional zone were structurally distinct from those in the mouse suggesting that they are unique biologically. Moreover, the trophoblasts in the bat junctional zone have histological differences within the layer itself, with marked differences in the cells abutting the labyrinth or decidua, indicating there may be multiple trophoblast subtypes within this layer (**Figure 1B, 1E**). Periodic Acid–Schiff (PAS) staining was used to detect glycogen in mouse and bat placental sections, enabling comparison of junctional zone organization and metabolic specialization across species. PAS staining was observed in junctional trophoblasts adjacent to the labyrinth zone and in some decidual cells of the bat placenta. Similarly, faint staining was detected in glycogen trophoblasts and decidua of the mouse placenta (**Figure S1C**, **S1D**). While these findings suggest potential functional similarities in glycogen storage between species, it remains unclear whether the extent of PAS staining in bats is influenced by gestational stage. Endothelial and fibroblast populations within the labyrinth were visualized by IHC using an antibody against Vimentin. Vimentin staining in the labyrinth of the bat resembled that of the mouse, where it stained fetal microvasculature. (**Figure 1A, 1C, 1D, 1F**, bottom panels and **Figure 1B, 1E** right panels). The lack of Vimentin staining also clearly defined the junctional zones in both the bat and mouse placentas. Viewed in conjunction, cytokeratin and Vimentin staining also clearly delineated two separate blood supplies in both the bat and mouse placentas, with narrow layers of fetal endothelial vasculature and placental syncytia separating the two blood supplies (**Figure 1C, 1F**). In mice, rats, and most other rodents there are two distinct layers of fused syncytiotrophoblast layers that surround the maternal blood supply to separate it from fetal blood. In contrast, the Jfb placentas were hemomonochorial, characterized by a single layer of fused syncytiotrophoblast separating maternal blood from fetal tissues. This architecture aligns with previous reports in other microbat lineages, including phyllostomid bats such as vampire bats^15^. However, comparative data across insectivorous and Old World microbat species remain limited, and further investigation is needed to assess the evolutionary breadth of this feature. The presence of morphologically distinct syncytial trophoblasts supports an invasive placental architecture in Jfb, marked by direct contact between maternal blood and the syncytiotrophoblast layer.

**Figure 1.**
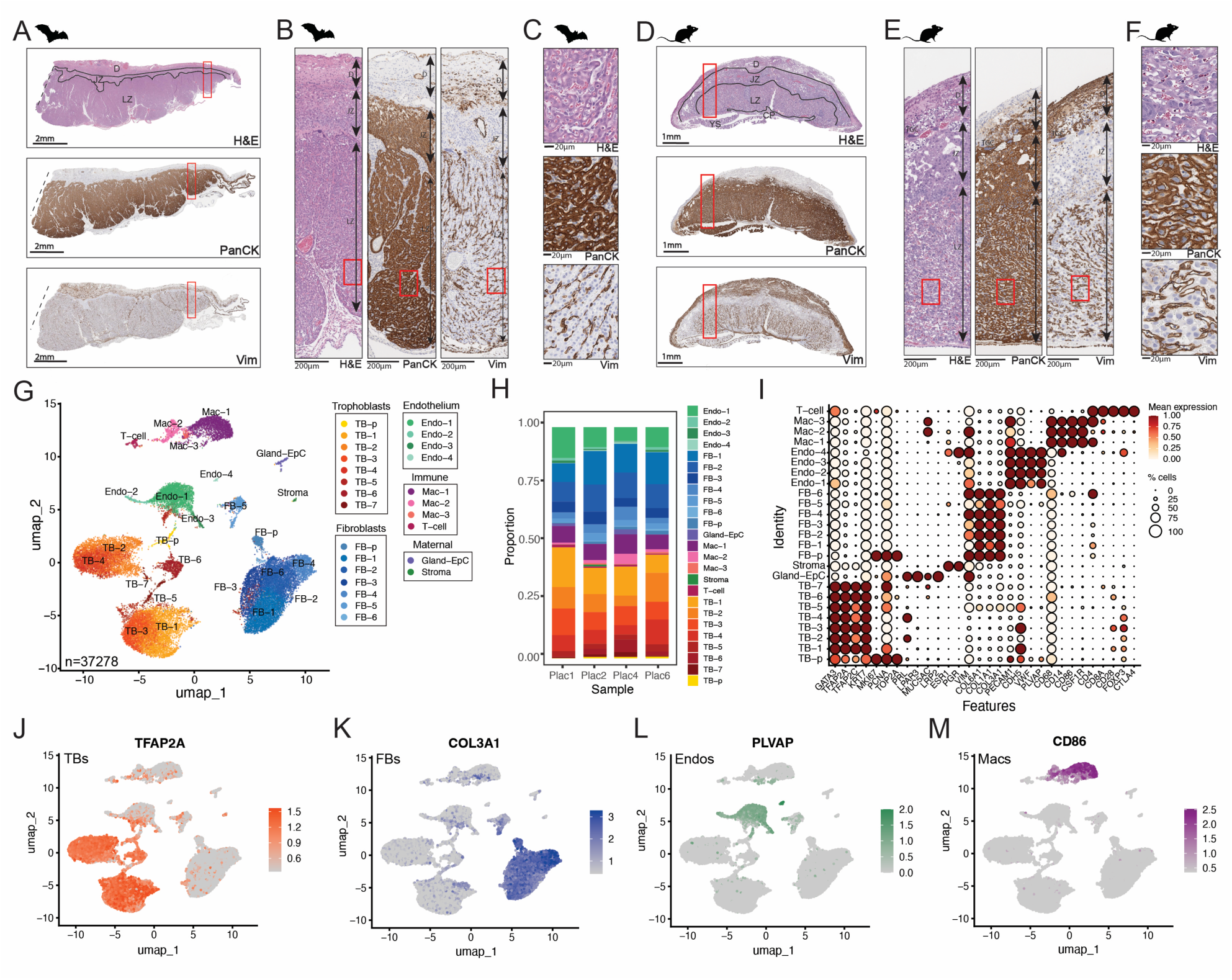
Cellular Landscape of the Jamaican Fruit Bat Placenta Revealed by Histology, Immunohistochemistry, and snRNA-Seq. **(A-F)** Representative histological and immunohistochemical staining of the Jamaican fruit bat (Jfb) placenta (A–C) and mouse placenta (D-F) at comparable gestational stages. (A, D) Low-magnification images show hematoxylin and eosin (H&E) staining (top row), pan-cytokeratin (Pan-CK, middle row) immunohistochemistry, and vimentin (Vim, bottom row) immunohistochemistry. Placental regions, including decidua (D), labyrinth zone (LZ), junctional zone (JZ), yolk sac (YS) labeled and regions are separated with black lines in (A) and (D). (B, E) Cross-sectional zoomed views of H&E (top), Pan-CK (middle), and Vim (bottom) staining highlight structural features within the placental layers. (C, F) High-magnification images show detailed morphology in H&E (top), Pan-CK (middle), and Vim (bottom) stained sections. Scale bars are shown at the bottom of each image. **(G)** UMAP visualization of single-nucleus RNA-seq data from four Jamaican fruit bat (Jfb) placentas, integrating four unique samples. A total of 37,278 high-quality nuclei were analyzed following SCTransform normalization and Harmony integration. **(H)** Proportional contribution of each sample to the 25 identified transcriptional clusters, demonstrating consistent cluster representation across distinct placentas. **(I)** DotPlot of canonical marker gene expression used to assign cluster identities, including markers for trophoblasts, fibroblasts, endothelial cells, macrophages, and T-cells. Scale at right. **(J–M)** FeaturePlots showing restricted expression of representative markers for trophoblasts (J, TFAP2C), fibroblasts (K, COL3A1), endothelial cells (L, PLVAP), and macrophages (M, CD86). Scale at right.

### Single-nucleus transcriptomic profiling of the Jfb placenta

Building on these histological findings, we next sought to define the cellular and transcriptional landscape of the Jfb placenta using single-nucleus RNA sequencing (snRNA-seq). This approach was chosen based on previous findings that snRNA-seq is necessary to capture the syncytiotrophoblast in human placental tissue^19,20^, which is often underrepresented in single-cell datasets due to its large, multinucleated structure and fragility during dissociation. Nuclei were isolated from four placentas, including one near full-term (Plac1) and three from mid-to-late gestation stages (Plac2, Plac4, and Plac6). Following quality control and filtering, we obtained a total of 37,278 high-quality nuclei for downstream analyses. This analysis resolved 25 transcriptionally distinct clusters across all samples, which were present in samples at near-equivalent ratios and expressed distinct cluster-associated genes (**Figure 1G, 1H, S2A, S2B, Table S1**). There were no significant differences between the late gestation Plac1 sample and those from tissues collected earlier in gestation. Despite efforts to assign fetal sex based on transcriptomic data, we were unable to do so due to the lack of annotated Y-linked genes in the *Artibeus jamaicensis* genome. Annotation attempts using Y-linked gene sequences from closely related bat species also failed to identify homologous loci in the available Jfb reference. To assign cell type identities to the 25 transcriptional clusters, we used a curated panel of canonical marker genes based on well-characterized human placental and immune cell populations^19,21–24^ (**Figure 1I**). Although the data were derived from Jfb tissue, these conserved markers provided a robust framework for interpreting bat placental cell types. Trophoblast clusters were identified by expression of genes such as *GATA3*, *TFAP2A/C*, and *KRT7*. Stromal and fibroblast populations were distinguished by expression of *ESR1*, *PGR*, as well as extracellular matrix genes including *VIM*, *COL6A1*, *COL1A1*, and *COL3A1*. Actively cycling cells were marked by proliferation-associated genes such as *MKI67*, *TOP2A*, and *PCNA*. Endothelial cells expressed *PECAM1*, *VWF*, *CDH5*, and *PLVAP*, while macrophage populations were identified by expression of *CD68*, *CD14*, *CD86*, and *CSF1R*. T-cell populations were annotated based on expression of *CD4*, *CD8A*, *CD28*, *FOXP3*, and *PDCD1*. A subset of clusters was also characterized by expression of decidual gland-associated markers including *PRL*, *LPAR3*, *MUC5AC*, *LRP2*, and *CDH2*, consistent with secretory epithelial cell identity. The presence of decidual glands was supported by H&E staining, which revealed glandular structures in placentas with intact uterine sections (**Figure S1F**).

Using the panel of canonical marker genes described above, we assigned identities to transcriptional clusters corresponding to major placental cell types. Trophoblast clusters were labeled as TB-p (proliferating) and TB-1-7, representing a spectrum of trophoblast subtypes likely reflecting progenitor, differentiating, and specialized states (**Figure 1G**). Fibroblast populations were annotated as FB-p (proliferating) and FB-1-6, capturing heterogeneity in stromal and mesenchymal populations. Four endothelial clusters (Endo-1-4) were identified based on vascular markers, while three macrophage clusters (Mac-1-3) reflected immune cell diversity within the placental microenvironment (**Figure 1G**). Lastly, a T-cell cluster was identified based on expression of T lymphocyte markers. FeaturePlots further demonstrated the specificity and restricted expression patterns of key marker genes across annotated clusters (**Figure 1J-1M, S2C-E**).

### Identification of Bat Trophoblast Markers and Pregnancy-Specific Glycoproteins

Given the limited understanding of trophoblast biology in bats, we sought to define the molecular signatures of bat trophoblasts at single-nuclei resolution. Building on our initial marker-based cluster annotations, we classified cell populations as either trophoblast or non-trophoblast (**Figure S2F**) and performed differential gene expression analysis using the MAST framework. This approach enabled the identification of genes significantly enriched in trophoblast clusters relative to all other cell types, expanding our view of trophoblast identity beyond canonical markers derived from human studies. We identified approximately 3,700 trophoblast-enriched genes (coding and non-coding) using a log₂ fold change threshold of 2 and an adjusted p-value cutoff of 0.05 (**Table S2**). However, due to redundancy in the Jfb genome annotation, including multiple LOC entries for the same locus, the true number of unique trophoblast genes is likely lower. This gene set included both known placental regulators and several uncharacterized or poorly annotated loci (**Figure 2A, 2B**). For example, *SHANK2*, a scaffolding protein associated with synaptic architecture, may reflect structural or signaling functions in trophoblasts. *CHRDL2*, a BMP antagonist, is potentially involved in modulating trophoblast differentiation. Additional candidates such as *CDH8* (a cadherin), *TRIM6* and *TRIML2* (ubiquitin ligases), and *SAXO1* (a microtubule stabilizer) suggest unique cytoskeletal, signaling, or adhesion-related roles. Other enriched transcripts included solute carriers (*SLC9A2*, *SLC9C2*), transcription factors (*RFX4*), and endogenous retroviral elements such as *LOC128627411*, annotated as a group S71 envelope polyprotein-like gene. Together, these findings define a broad repertoire of bat-specific trophoblast genes and provide a foundation for understanding species-specific placental adaptations.

**Figure 2.**
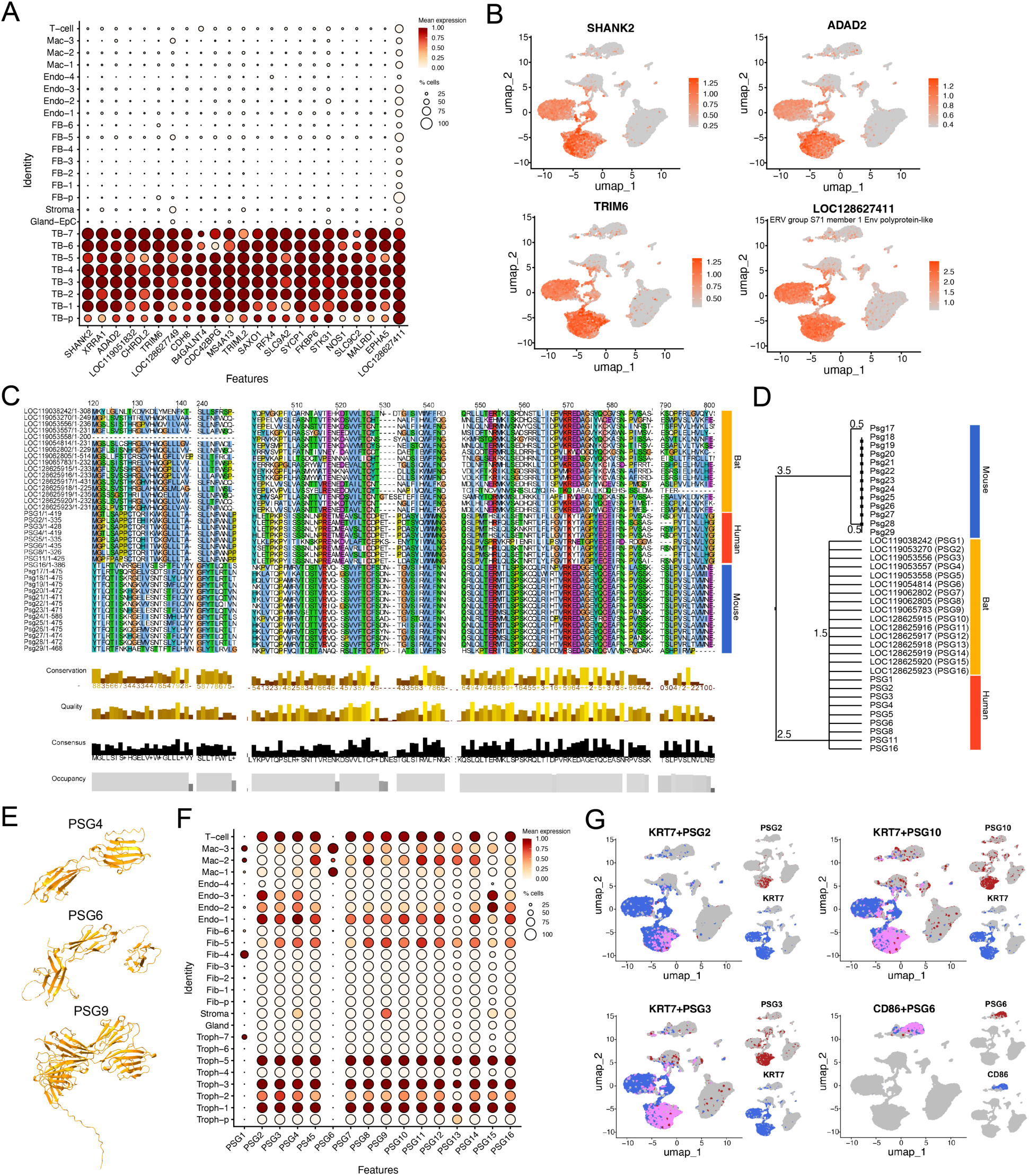
Identification of Bat Trophoblast Markers and Pregnancy-Specific Glycoproteins (PSGs). **(A)** DotPlot of genes enriched in Jfb placental trophoblasts versus non-trophoblasts as determined by differential expression analysis. Key at right. **(B)** FeaturePlots of select genes identified in (A). Scale at right. **(C)** Conserved regions across full PSG alignments of candidate bat PSGs (yellow-orange bar) and select human PSGs (red bar) or mouse Psgs (blue bar) using MEGA11. **(D)** Phylogenetic tree based on ClustalW multiple sequence alignment of candidate bat PSGs with human and mouse PSG families, showing closer homology of bat PSGs to human than to mouse sequences. **(E)** Domain architecture of representative bat PSG proteins, predicted by AlphaFold3, demonstrating conserved Ig-like domain structures typical of PSGs. **(F)** DotPlot of candidate bat PSG gene expression across placental cell types, demonstrating enrichment in trophoblast populations. **(G)** FeaturePlots showing the co-expression of the indicated bat PSGs (in red) with KRT7 (blue), a trophoblast marker, supporting their localization in trophoblast populations or the unique expression of PSG6 in a macrophage cluster localized with CD86 (blue, bottom right).

Pregnancy-specific glycoproteins (PSGs) represent one of t. he most prominent and lineage-restricted gene families expressed by trophoblasts in several mammalian species. In humans, PSGs are secreted by the syncytiotrophoblast and play key roles in modulating maternal immune responses and supporting fetal development^25–30^. Despite their well-characterized roles in primates and rodents, PSGs have not been extensively characterized in bats. Given their status as a defining feature of trophoblast gene expression in other mammals, we hypothesized that bats may also express PSG-like genes as part of their placental program. To explore this possibility, we surveyed the *Artibeus jamaicensis* genome for both annotated CEACAM family members and uncharacterized CEACAM-like genes listed as LOC entries. This search yielded 31 candidate PSG-like genes (**Table S3**). Given that true PSGs are distinguished by the absence of transmembrane domains, we applied DeepTMHMM^31^ to predict protein topology, identifying 24 candidates lacking transmembrane regions, consistent with a possible soluble PSG identity.

To evaluate orthology and evolutionary relationships, we aligned these 24 bat PSG candidates with known human PSGs (PSG1–8, 11, 16) and mouse Psgs (Psg17–29) using ClustalW. Among these 24 bat PSG candidates, 16 genes shared high sequence consensus with human and mouse PSGs. We visualized conserved regions among PSGs across species with Jalview (v2.11.4)^32^ (**Figure 2C**). Phylogenetic analysis in MEGA11^33^ showed that bat PSGs cluster more closely with human than with mouse PSGs, suggesting evolutionary conservation with the primate lineage (**Figure 2D**). We then predicted 3D structures for these 16 candidates using AlphaFold 3.0^34^, alongside representative human and mouse PSGs. Structural overlays revealed conservation of immunoglobulin-like domains and domain organization, supporting their classification as bona fide PSGs (**Movie S1**). All 16 high-confidence bat PSGs encoded two to three extracellular Ig-like domains, including an N-terminal IgV domain followed by one or more IgC domains, an architecture characteristic of PSGs across mammals (**Figure 2E**). Based on this integrated analysis of domain structure, sequence conservation, and structural homology, we designated these genes as *PSG1–16*, representing a previously unrecognized expansion of the PSG family in bats (**Table S3**).

To validate their functional relevance, we assessed the expression of bat PSGs in the full snRNA-seq dataset. Most PSGs were robustly expressed in trophoblast clusters, as assessed by their strong colocalization with *KRT7* (**Figure 2F, 2G**). In contrast, PSG6 displayed a distinct expression pattern restricted to a macrophage population (**Figure 2G**), suggesting that some bat PSGs may have evolved noncanonical roles outside the trophoblast lineage, potentially contributing to immunoregulatory functions at the maternal-fetal interface.

### Trophoblast diversity and specialization in the Jfb placenta

To gain deeper insights into the diversity and specialization of bat trophoblasts, we focused our analysis on the subset of clusters identified as trophoblasts. This allowed us to resolve distinct transcriptional states and explore putative lineage trajectories within trophoblasts comprising the Jfb placenta. To accomplish this, we subsetted all trophoblast populations described above based on expression of trophoblast-enriched genes and re-analyzed these clusters to improve resolution of trophoblast cell types. This analysis retained seven distinct trophoblast (TB) clusters. TB 1–2 formed a separate axis on the UMAP, while TB 4–6 clustered along another (**Figure 3A**). TB-3 emerged as a uniquely distinct population, and a proliferating trophoblast cluster (TB-p) was also preserved from the original analysis (**Figure 3A**). The ratios of these cell populations were near-equivalent between samples (**Figure 3B, S3A**). Each of these distinct groups of clusters expressed distinct markers (**Figure S4B**). To better characterize the molecular identity of each trophoblast population, we examined gene expression profiles across clusters (**Table S4**). TB-1–2 were enriched for genes associated with signaling, transcriptional regulation, and WNT pathway activity, including *CHST4*, *SALL3*, *WNT3A*, and *OTX1* (**Figure S3B**). GO terms in these clusters reflected roles in cadherin binding, chromatin remodeling (e.g., histone binding, ATP-dependent DNA activity), and translational control, suggesting a progenitor-like, transcriptionally dynamic state (**Figure S3C**). TB-3 stood out as a transcriptionally distinct population marked by genes involved in neuroendocrine signaling, immune modulation, and cell adhesion, such as *PTPRN*, *CCR7*, *NFATC1*, and *GDNF* (**Figure S3B, S3C**). Corresponding GO terms included GTPase binding, actin binding, and ubiquitin-like protein transferase activity, supporting a unique regulatory identity for this cluster (**Figure S3C**). In contrast, TB-4–6 expressed genes linked to metabolic processes, epithelial polarity, and hormone responsiveness, including *CCKAR*, *GRHL3*, *SLC2A4*, and *PLA2G4D* (**Figure S3B**). GO enrichment in these clusters highlighted functions such as structural constituents of ribosomes, SMAD binding, growth factor interactions, and unfolded protein binding, indicative of syncytial specialization and metabolic activation (**Figure S3C**). These distinct gene expression signatures suggest functional specialization among trophoblast subsets.

**Figure 3.**
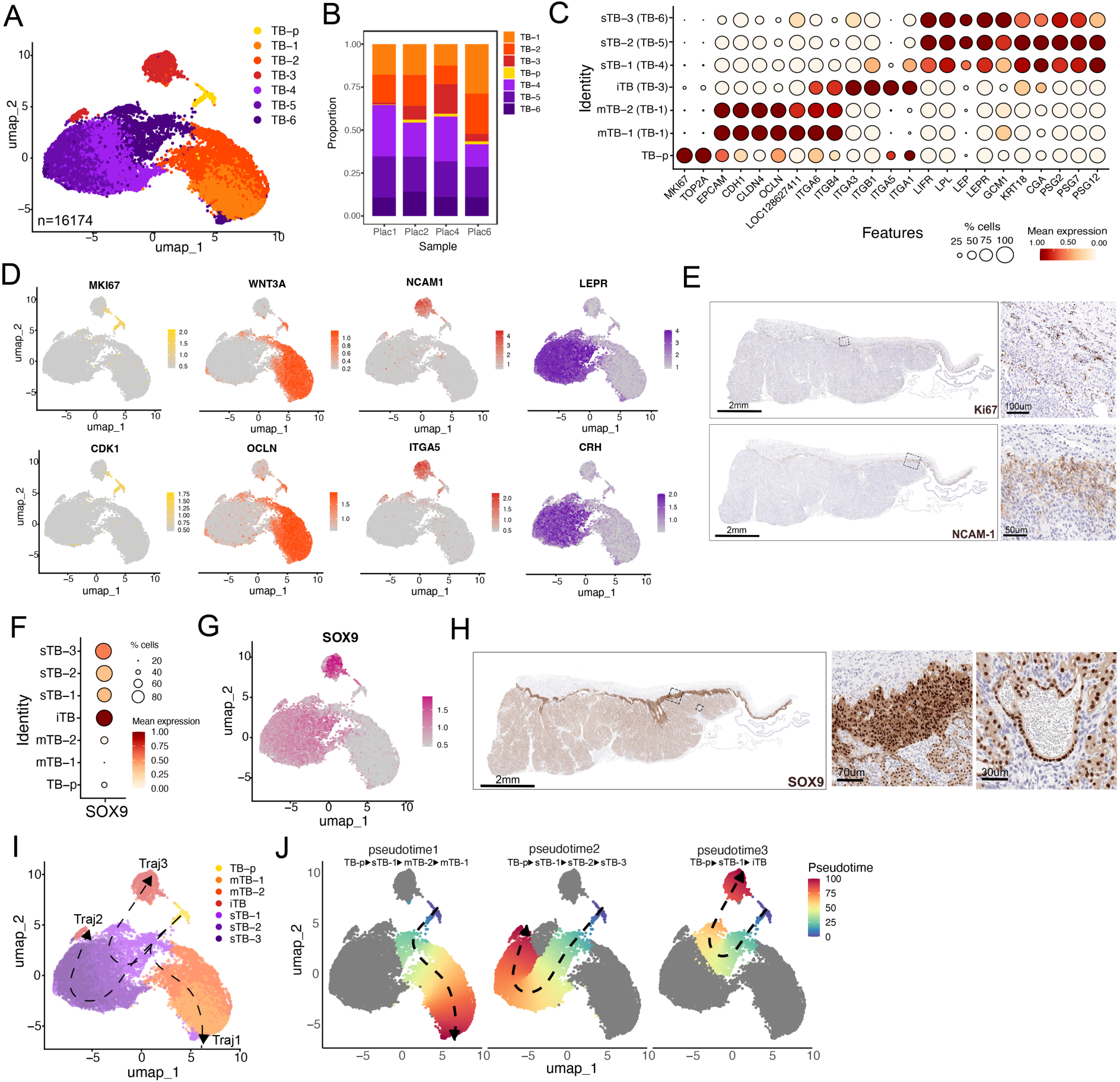
Transcriptional Heterogeneity and Differentiation Trajectories of Trophoblast Populations in the Jamaican Fruit Bat Placenta. **(A)** UMAP visualization of subsetted trophoblast populations from single-nucleus RNA-seq data, resolving seven distinct clusters. **(B)** Sample contribution to each trophoblast cluster, showing comparable representation across placental samples. **(C)** DotPlot of canonical human trophoblast subtype markers (CTB, EVT, STB) and select bat PSGs across bat trophoblast clusters resolving the clusters identities in (A) as follows: TB-p (TB-p) (proliferating trophoblasts), mTB-1 and mTB-2 (TB-1 and TB-2) (mononuclear trophoblasts), iTB (TB-3) (invasive trophoblasts), and sTB (sTB-1, sTB-2, and sTB-3) (syncytial trophoblasts). **(D)** FeaturePlots of key markers associated with proliferative (*MKI67*, *CDK1*), epithelial (*WNT3A*, *OCLN*), invasive (*NCAM1*, *ITGA5*), and syncytial (*LEPR*, *CRH)* states. **(E)** Immunohistochemistry for Ki67 and NCAM1 on Jfb placental sections, validating proliferative and invasive trophoblast populations, respectively. **(F–G)** SOX9 expression across clusters by DotPlot (F) and FeaturePlot (G), showing strong expression in iTB and moderate expression in syncytial sTB populations. **(H)** Immunohistochemistry for SOX9 in bat placental tissue. Images at right are zoomed from black boxes and show SOX9 localization in iTB and sTB populations. Scale at bottom. **(I)** Slingshot trajectory analysis of trophoblast populations, using the proliferating trophoblast cluster (TB-p) as the root node. Three major lineage trajectories were identified: one leading toward mTB, one toward sTB, and one toward iTB. **(J)** UMAP visualization of trophoblast differentiation trajectories, colored by pseudotime (rainbow scale) and with arrows indicating directionality; red shading denotes terminal differentiation states.

Next, we assessed the expression of canonical markers associated with known human trophoblast subtypes, including cytotrophoblasts (CTBs), extravillous trophoblasts (EVTs), and syncytiotrophoblast (STB), to annotate trophoblast populations in the Jfb placenta^19,21–24^. These included proliferation and epithelial markers such as *MKI67*, *TOP2A*, *EPCAM*, *CDH1*, *TJP1*, and *CLDN4*; EVT-associated integrins and immune modulators like *ITGA5*, *ITGA1*, *ITGB1*, *NCAM1*, *LGALS3*, and *MMP14*; and syncytial markers including *CGA*, *LEP*, *LIFR*, *CRH*, *GDF15,* and various *PSGs* (**Figure 3C, 3D**). TB-1–2 expressed high levels of epithelial and progenitor-associated CTB markers, consistent with a mononuclear, less differentiated state. TB-3 showed strong expression of invasive and immunomodulatory genes, including multiple integrins and *NCAM1*, as well as elevated *SOX9*, suggesting an EVT-like identity with a transcriptional program associated with invasion and intermediate differentiation (**Figure 3C, 3F, 3G**, **S3B**). TB-4–6 expressed a range of syncytial and hormone-related markers, including moderate levels of *SOX9*, supporting their classification as differentiated, hormone-active STB populations. Based on these integrated transcriptomic and spatial patterns, we re-annotated the clusters as follows: TB-1–2 as mononuclear trophoblasts (mTB), TB-3 as invasive trophoblasts (iTB), and TB-4–6 as syncytial trophoblasts (sTB). To validate these transcriptional profiles *in situ*, we performed IHC for Ki67, NCAM1, and SOX9. Ki67 marked proliferative trophoblasts, primarily localized to the junctional zone (**Figure 3E**, top), while NCAM1 expression was restricted to cells at the maternal-fetal interface (**Figure 3E**, bottom), consistent with an invasive phenotype. SOX9 protein was most abundant in iTB populations, with moderate expression in sTB and little to no expression in mTB or TB-p (**Figure 3H**). Together, these findings establish a transcriptional framework for trophoblast lineage specialization in the Jfb placenta.

To infer differentiation dynamics among bat trophoblast populations, we applied Slingshot trajectory analysis using the proliferating trophoblast cluster (TB-p) as the root. This approach revealed three distinct developmental trajectories originating from TB-p, reflecting branching fates within the trophoblast lineage (**Figure 3I, 3J**). The first trajectory extended from TB-p to sTB-1, then progressed to mTB-2 and ultimately to mTB-1. This path suggests a lineage originating in a proliferative, undifferentiated state and culminating in mTBs, potentially representing a progenitor-to-CTB-like maturation sequence. The second trajectory also initiated in TB-p and sTB-1 but instead progressed through sTB-2 and terminated in sTB-3. This pattern indicates a committed syncytial differentiation pathway, with sTB-3 representing a terminal, mature syncytiotrophoblast state. The third trajectory led from TB-p to sTB-1 and then diverged toward iTB, the invasive trophoblast cluster. This suggests that invasive trophoblasts may arise via a shared intermediate state with syncytial lineages before diverging toward an EVT-like fate. To define gene expression dynamics along these lineages, we used fitGAM to model gene expression changes across pseudotimes. This analysis revealed trajectory-specific transcriptional programs. For instance, LEPR expression increased along the syncytial (sTB) trajectory, consistent with its role in hormone signaling in differentiated trophoblasts, while NCAM1 was specifically upregulated along the invasive (iTB) trajectory, reflecting its association with migratory, EVT-like states (**Figure S3D**). Together, these findings support a model in which sTB-1 serves as a key intermediate state from which multiple trophoblast fates arise, including terminal syncytial, mononuclear, and invasive lineages.

### Mapping immune cell populations in the Jfb placenta

We identified three transcriptionally distinct macrophage populations and one T-cell population in the Jfb placenta by snRNA-seq (**Figure 4A, 4B**, **S4A**, **Table S5**). All immune populations expressed canonical markers supporting their identity and functional relevance in the placental environment (**Figure 4C**). Macrophage populations expressed markers such as *CD68*, *MARCO*, *CD14*, *CD163*, *MRC1*, and *CSF1R*, whereas the T-cell population expressed *CD4*, *CD8A*, *IL7R*, and *FOXP3*. Marker expression revealed notable heterogeneity among macrophage populations: for example, CD14 was enriched in one cluster and CD163 in another, suggesting functionally specialized subsets (**Figure 4C, 4D**, **S4B**). In contrast, the single T-cell population exhibited a mixed transcriptional profile, expressing genes associated with naïve T-cells (*SELL*, *TCF7*, *CCR7*), effector function (*GZMB*, *IFNG*, *PRF1*), and regulatory T-cells (*FOXP3*, *IL2RA*, *IKZF2*) (**Figure S4B**). Broad expression of TCR signaling and activation markers (*CD3D*, *ZAP70*, *LCK*) confirmed its identity as a bona fide T-cell population. To validate and localize immune cells *in situ*, we performed IHC for CD45 (pan-leukocyte), CD3ε (T-cells), CD68, and F4/80 (a marker of tissue-resident macrophages in mice). CD45⁺ cells were enriched in the junctional zone, particularly surrounding the microvasculature (**Figure 4E**). CD3ε⁺ T-cells were rare but present in both the junctional zone and labyrinth. CD68⁺ macrophages were abundant in the junctional zone, and F4/80 showed strong, restricted staining of cells lining this region, consistent with a tissue-resident macrophage population (**Figure 4E**).

**Figure 4:**
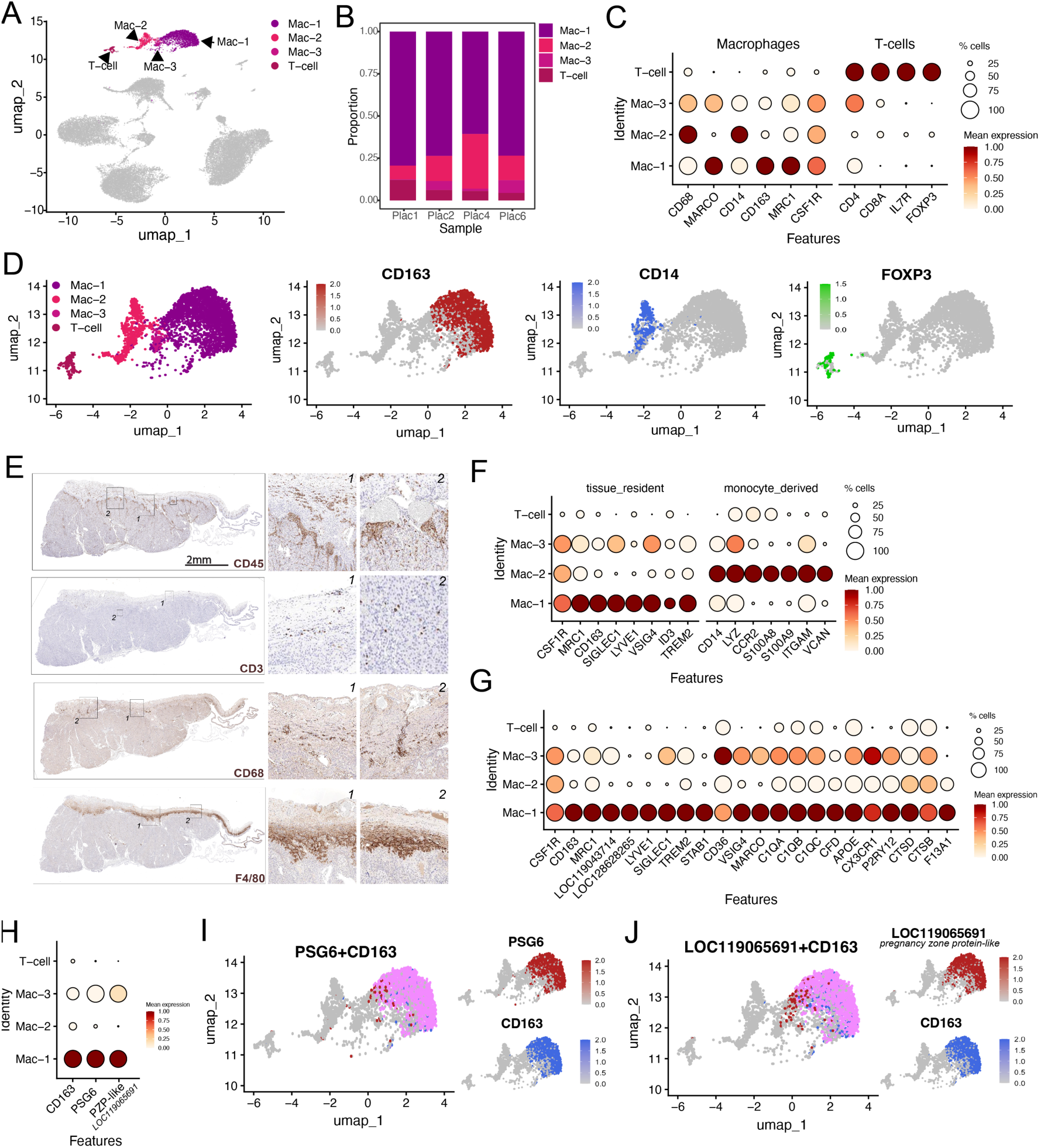
Immune Cell Diversity and Functional Specialization at the Maternal-Fetal Interface of the Jamaican Fruit Bat Placenta. **(A)** UMAP visualization of immune cell populations identified in single-nucleus RNA-seq data from the Jamaican fruit bat placenta, resolving three macrophage clusters (Mac-1, Mac-2, Mac-3) and one T-cell cluster. **(B)** Proportional contribution of each sample to immune clusters, demonstrating consistent representation across independent placentas. **(C)** DotPlot of canonical macrophage and T-cell markers, including *CD68*, *MARCO*, *CD14*, *CD163*, *MRC1*, *CSF1R* (macrophages) and *CD4*, *CD8A*, *IL7R*, *FOXP3* (T-cells). **(D)** FeaturePlots of representative immune markers *CD163* (red), *CD14* (blue) and *FOXP3* (green) shown on a UMAP subsetted to dimensions capturing only immune cell populations (left), improving resolution of cluster-specific expression patterns. **(E)** Immunohistochemistry for CD45 (pan-leukocyte marker), CD3 (T-cells marker), CD68, and F4/80 (macrophage markers) showing localization of immune cells predominantly to the junctional zone, with rare T-cells present in both junctional and labyrinthine regions. Scale bar, 2 mm. **(F)** Expression of tissue-resident versus monocyte-derived macrophage markers across macrophage clusters by DotPlot. **(G)** Expression of Hofbauer cell–associated genes demonstrating that Mac-1 strongly resembles human placental Hofbauer cells. **(H)** DotPlot showing differential enrichment of *LOC119065691* (a pregnancy zone protein–like homolog, PZP) and *PSG6* in Mac-1 relative to other macrophage populations or T-cells. **(I, J)** Co-expression of (I) *PSG6* (red) or (J) *LOC119065691/PZP-like* (red) and *CD163* (blue) within Mac-1 as assessed by FeaturePlot/

To determine whether the macrophage populations in the bat placenta were tissue-resident or derived from recruited monocytes, we examined expression of canonical marker genes. We used a curated panel that included markers of tissue-resident macrophages (e.g., *CSF1R*, *MRC1*, *CD163*, *SIGLEC1*, *LYVE1*, *VSIG4*, *ID2*, *TREM2*) and monocyte-derived macrophages (e.g., *CD14*, *FCGR3A*, *LYZ*, *CCR2*, *S100A8*, *ITGAM*, *VCAN*). This analysis revealed that the Mac-1 population strongly expressed tissue-resident markers (**Figure 4F**). Mac-3 showed lower but detectable expression of these same markers (**Figure 4F**). In contrast, Mac-2 was enriched for monocyte-derived macrophage markers, suggesting it represents a more inflammatory, recruited population (**Figure 4F**). Given that Mac-1 displayed strong expression of tissue-resident macrophage markers, we next determined whether this population resembled Hofbauer cells, specialized fetal macrophages found in the human placenta. To assess this, we examined expression of a curated set of Hofbauer cell–associated genes, including core macrophage markers (*CSF1R*, *CD163*, *MRC1*, *LYVE1*, *SIGLEC1*), scavenger receptors (*STAB1*, *CD36*, *VSIG4*), complement components (*C1QA*, *C1QB*, *C1QC*), and lipid handling genes (*APOE*), as well as microglia-like genes (*CX3CR1*, *P2RY12*)^21^. Mac-1 expressed nearly all these markers at high levels, consistent with a Hofbauer-like identity (**Figure 4G**). In contrast, Mac-2 and Mac-3 lacked broad expression of these genes, supporting the notion that Mac-1 represents a fetal-derived, tissue-resident macrophage population, while other subsets may reflect distinct or more inflammatory macrophage lineages.

Because the Mac-1 cluster expressed a broad range of Hofbauer-like markers, we next sought to identify additional genes that might clarify its identity and function. Using gene marker enrichment analysis, we found that this cluster was specifically enriched for *LOC119065691* (log_2_FC= 5.86, p_adj_= 1.84E-260), which encodes a pregnancy zone protein-like (PZP) homolog (**Figure 4H, Table S5**). This gene was expressed almost exclusively in the Mac-1 population and showed strong co-expression with *CD163*, a hallmark of tissue-resident, anti-inflammatory macrophages (**Figure 4I**). PZP is a high-molecular-weight protein elevated during pregnancy in humans and mice and has been proposed to play roles in protein stabilization^35,36^. *PSG6*, one of the 16 bat pregnancy-specific glycoproteins, also showed strong, selective expression in this same macrophage cluster (**Figure 4H, 4I**), suggesting a potential immunomodulatory role distinct from other PSGs expressed in trophoblasts. Together, these findings suggest that Mac-1 represents a Hofbauer-like, fetal-derived macrophage population with a unique transcriptional signature marked by high *PZP* and *PSG6* expression.

### Cross-species Analysis of Bat, Human, and Mouse Placental Cell Types

To place the cellular landscape of the Jfb placenta in an evolutionary framework, we next performed cross-species integration of our bat snRNA-seq dataset with publicly available single-nucleus RNA-seq data from first-trimester human and mid-gestation (E12) mouse placentas (**Figure 5A**)^20,37,38^. Each dataset contained the expected placental cell types, including trophoblasts, endothelial cells, fibroblasts, and immune cells, based on canonical marker expression (**Figure S5A-S5D**)^19^. To enable direct comparisons, datasets were quality-controlled separately and restricted to a shared set of 13,546 genes. We standardized gene nomenclature by converting mouse gene symbols to uppercase, aligning with human and bat annotations. Following integration and clustering, we restored species-specific annotations, including the lowercase formatting of mouse genes. This workflow preserved both cross-species alignment and the ability to detect species-specific transcriptional features. The final integrated object contained 110,613 nuclei: 37,278 from bat, 22,753 from human, and 50,582 from mouse, and resolved into 22 transcriptionally distinct clusters (**Figure S5F**).

**Figure 5.**
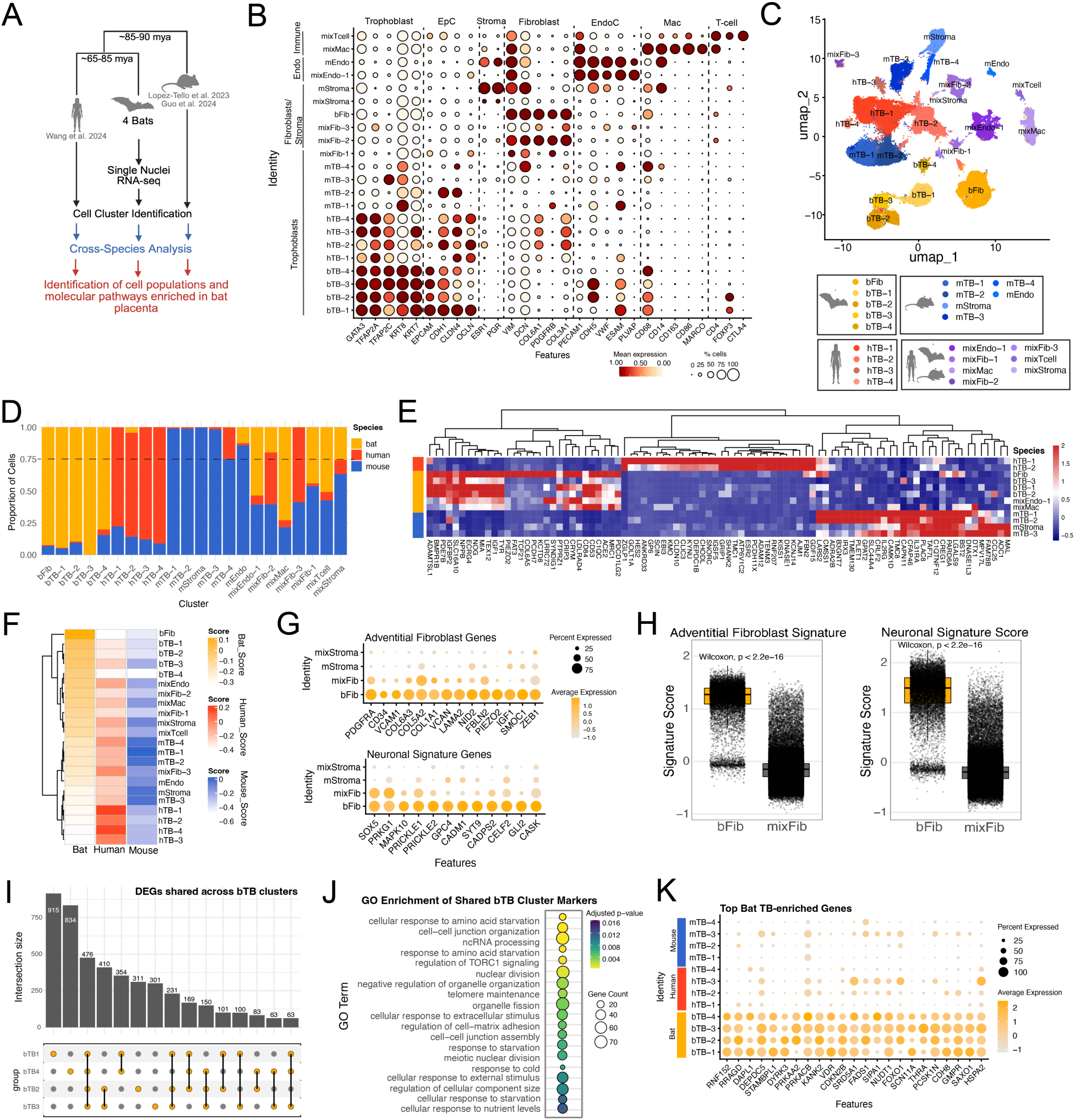
Single-Nucleus RNA Sequencing Defines Conserved and Species-Specific Placental Features Across Bats, Humans, and Mice. **(A)** Schematic of cross-species single-nucleus RNA-seq integration, comparing Jamaican fruit bat (Jfb), human, and mouse placental datasets. mya, million years ago **(B)** DotPlot showing expression of canonical marker genes across integrated clusters, highlighting trophoblast, fibroblast, endothelial, stromal, and immune populations. Scale at bottom. **(C)** UMAP visualization of the integrated dataset, identifying species-specific and mixed-species clusters. Bat-derived clusters are shown in yellow-orange and designated with a "b" prefix, human clusters are shown in red and designated with an "h," and mouse clusters are shown in blue and designated with an "m." Mixed-species clusters are shown in purple and labeled as "mix." **(D)** Proportional contribution of bat, human, and mouse nuclei across clusters, revealing strong species specificity among trophoblasts and greater conservation among non-trophoblast lineages. **(E)** Contribution of species-restricted gene signatures across major placental cell types as shown by heatmap. Key at right. **(F)** Heatmaps show the scaled expression (Z-score) of bat, human, and mouse gene signature scores across all annotated clusters. Signature scores were calculated per cell based on the top 1,000 differentially expressed genes for each species, identified using MAST (FDR < 0.05, log₂FC > 0.25), and then averaged by cluster. Each column represents a species-specific gene signature, and each row represents a defined placental cluster as indicated. Row names indicate cluster identity, and color intensity reflects relative enrichment within each cluster. Cluster order was maintained across species. Bat is shown in yellow-orange, human in red, and mouse in blue. **(G)** DotPlots of select genes associated with adventitial (top) and neuronal (bottom) gene sets in bFib versus mixFib and stromal popualtions. Key at right. **(H)** Boxplots show signature scores for curated adventitial (left) and neuronal (right) gene sets in bat-specific fibroblasts (bFib) versus all mixed-species fibroblasts (mixFib). bFib cells showed significantly higher enrichment for both signatures (as assessed by Wilcoxon test)**. (I)** ComplexUpset plot showing shared and unique DEGs among bTB-1, bTB-2, bTB-3, and bTB-4 clusters. Each vertical bar represents the number of DEGs associated with a specific combination of subpopulations (highlighted by connected dots below), while horizontal bars indicate the total number of DEGs per cluster. **(J)** GO pathway analysis of genes enriched in bTB clusters compared to human and mouse trophoblasts identified significant enrichment for pathways related to nutrient sensing, autophagy, and environmental stress responses. Dot size represents the number of genes associated with each term, and color indicates statistical significance (adjusted p-value). **(K)** DotPlots of select bTB-enriched genes associated with pathways shown in J across all TB clusters. Key at right.

To assign species identity, we labeled clusters based on their cellular composition: clusters with >75% of nuclei from a single species were designated as bat (BAT), human (HUM), or mouse (MOU), while more heterogeneous clusters were labeled as mixed (MIX) (**Figure S5G**). This approach yielded five bat-specific clusters (BAT-1 to BAT-5), four human-specific (HUM-1 to HUM-4), six mouse-specific (MOU-1 to MOU-6), and seven mixed clusters (MIX-1 to MIX-7) (**Figure S5G, S5H**). Cell type annotation using canonical lineage markers revealed strong species specificity among trophoblast clusters, whereas non-trophoblast clusters, such as endothelial, stromal, and immune populations, more frequently displayed interspecies overlap (**Figure 5B, 5C**).

Among bat-specific clusters, four were enriched for trophoblasts (bTB-1 to bTB-4; BAT-2 to BAT-5), and one represented a fibroblast population (bFib; BAT-1). Human-specific clusters consisted exclusively of trophoblasts (hTB-1 to hTB-4; HUM-1 to HUM-4), reflecting the well-characterized trophoblast diversity of the early human placenta (**Figure 5C**). Mouse-specific clusters included four trophoblast groups (mTB-1 to mTB-4; MOU-1, −2, −4, −5), one stromal cluster (mStroma; MOU-3), and one endothelial cluster (mEndo; MOU-6). The mixed clusters (MIX-1 to MIX-7) were largely composed of non-trophoblast lineages, including endothelial cells (mixEndo-1; MIX-1), fibroblasts (mixFib-1-3; MIX-2, −4, and −5), macrophages (mixMac; MIX-3), stromal cells (mixStroma; MIX-7), and T-cells (mixT-cell; MIX-6).

We next quantified the species contribution to each cluster to assess the degree of interspecies overlap (**Figure 5D**). While some clusters, such as mixEndo-1 and mixMac, contained nuclei from all three species, others were highly species-restricted. For example, bTB-3 and bTB-4 were composed almost entirely of bat-derived nuclei (89.6% and 100%, respectively), and hTB-1 to hTB-4 were >77% human. All mouse-specific clusters were >98% mouse-derived. Among the mixed clusters, several displayed restricted diversity: mixFib-3 lacked bat nuclei, mixFib-1 was predominantly mouse and bat, and mixStroma was bat- and mouse-enriched with minimal human representation. These patterns suggest that non-trophoblast lineages are generally more conserved across species, while trophoblast populations are highly species-specific, likely reflecting divergent evolutionary pressures and developmental programs. A notable exception was the bat-specific fibroblast cluster, bFib (BAT-1), which contained ∼92% bat nuclei yet expressed conserved fibroblast markers such as *VIM* and *DCN* (**Figure 5B, 5D**).

To further define the species-specific transcriptional features underlying placental specialization, we performed differential expression analysis across all nuclei using MAST with a minimum expression threshold (min.pct = 0.25) and a log₂ fold change cutoff of 2. This analysis identified 2,013 bat-specific, 414 human-specific, and 77 mouse-specific genes (**Tables S6–S8**). Bat-enriched genes included those involved in growth factor signaling (*IGF1, NOG, BMPR1B*), immune regulation (*MRC1*), hormone activity (*ACE2, NPPB*), pigment biosynthesis (*TYR*), and extracellular signaling (*IGFBP2*). Human-specific genes included *KISS1*, *ADAM12*, and *KYNU*, while mouse-specific enrichments featured *TAF7L*, *PGR*, and *FAM135B*, suggesting divergent transcriptional programs even among conserved cell types.

To determine which cell populations contributed most to the species-enriched transcriptional programs, we calculated bat, human, and mouse signature scores for each cluster using the corresponding differentially expressed gene sets. These scores were visualized as a heatmap to highlight species-specific enrichment patterns across cell populations (**Figure 5F**). Mapping these results to annotated cluster identities revealed that human-enriched scores were highest in hTB-1 through hTB-4, mouse-enriched scores were prominent in mTB-1 through mTB-4 and mStroma, and bat-enriched scores were strongly elevated in bFib and bTB-1 through bTB-4. Mixed clusters including mixEndo-1, mixFib-1/2, mixMac, and mixTb exhibited lower and more variable enrichment, consistent with their heterogeneous composition. These results indicate that the most prominent bat-specific transcriptional signatures are concentrated within fibroblast and trophoblast populations.

To define the molecular features underlying the species-specific clustering of bFib cells, we next examined the gene expression programs that distinguished this population from other fibroblasts and stromal populations. In reviewing the bat-enriched gene list, we observed that many top markers were characteristic of adventitial fibroblasts, perivascular stromal cells found in large vessels that contribute to tissue remodeling, immune regulation, and mechanosensation. bFib cells selectively expressed canonical adventitial markers including *PIEZO2, PDGFRA, IGF1*, and *ZEB1*, as well as extracellular matrix components such as *VCAN, FBLN2, COL6A3, COL5A2, LAMA2, GPC4*, and *TSPAN5* (**Table S9**). Additional secreted regulators including *SMOC1, ADAMTS12*, and *NID2* supported an adventitial-like identity (**Figure 5G**). To validate this, we calculated an adventitial fibroblast signature score using a curated marker set^39,40^, which revealed significantly higher enrichment in bFib compared to other fibroblast populations (**Figure 5H**). bFib cells also expressed a number of genes associated with neuronal signaling and synaptic architecture, including *SOX5, PRKG1, MAPK10, PRICKLE1, PRICKLE2, GPC4, CADM1, SYT9, CADPS2, CELF2, GLI2*, and *CASK* (**Figure 5G**). A neuronal signature score based on this panel showed similarly strong enrichment in bFib cells relative to mixed-species fibroblasts (**Figure 5H**). This dual enrichment supports the presence of a transcriptionally distinct fibroblast subtype in the bat placenta that combines features of adventitial and neuron-associated fibroblasts.

Given that bat trophoblast clusters were among the strongest contributors to the bat-specific transcriptional signature, we next sought to identify the gene programs driving this enrichment. Using MAST, we performed differential expression analysis for each bat trophoblast population (bTB-1 through bTB-4) compared to human and mouse trophoblasts. We identified upregulated genes (log₂ fold-change > 2, adjusted p < 0.05) for each bat TB population. ComplexUpset analysis revealed that a substantial proportion of differentially expressed genes were shared across bTB-1, bTB-2, bTB-3, and bTB-4, highlighting a conserved core transcriptional signature among bat trophoblast subtypes (**Figure 5J**). Although most DEGs were broadly shared, bTB-1 and bTB-4 also exhibited unique subsets of enriched genes, suggesting additional lineage-specific specialization within proliferative and differentiated trophoblast states. Shared genes defined a conserved bat trophoblast signature, suggesting that although individual clusters exhibit distinct transcriptional profiles, they are unified by common molecular programs. To gain functional insight into these bat-enriched genes, we performed Gene Ontology enrichment analysis. This revealed several pathways specifically upregulated in bat trophoblasts, including cellular responses to amino acid starvation (*RNF152, RRAGD, DAPL1, DEPDC5*), regulation of TORC1 signaling (*RNF152, STAMBPL1, DYRK3, PRKAA2*), and responses to cold and metabolic stress. Additional enriched pathways included cell–cell junction organization (*CDH13, NLGN2, LIMS2, ANK2*) and non-coding RNA processing (*TDRD9, TERT, PIH1D2, FKBP6*), suggesting adaptations related to nutrient sensing, stress resilience, and specialized transcriptional control (**Figure 5K, 5L**). Together, these findings indicate that bat trophoblasts activate distinct molecular programs compared to their human and mouse counterparts, potentially reflecting species-specific strategies to support pregnancy under fluctuating environmental conditions and sustained metabolic demands.

### Derivation of Trophoblast and Decidua Organoids from the Jfb Placenta

The findings described above underscore the cellular complexity of the bat placenta and highlight the need for *in vitro* models that capture key trophoblast states. The presence of a high proportion of Ki67-positive cells in the Jfb placenta suggested the existence of actively cycling stem/progenitor populations. This proliferative signature indicated that it might be possible to isolate and expand these progenitors *in vitro* to generate organoids, as we have previously demonstrated in human, rhesus macaque, and pig placentas^41–43^. To test this, we sought to establish organoids representing both trophoblast and decidual gland epithelial lineages from the Jfb placenta. Labyrinth tissue, which is fetal-derived, was carefully separated from maternal-derived uterine and decidual tissue, followed by enzymatic digestion and 3D culture in Matrigel using a cocktail of growth factors previously optimized for cross-species organoid derivation^41–43^ (**Figure 6A**). This approach led to the successful generation of two distinct organoid types: trophoblast organoids (TOs) from labyrinth tissue and decidual organoids (DOs) from maternal decidua (**Figure 6B**). TOs and DOs were successfully established from all five Jfb placentas, confirming the feasibility and reproducibility of organoid generation from this species. Both TOs and DOs grew robustly in culture, with visible organoid structures emerging within 2 weeks post-isolation (**Figure S6A**). Organoid cultures required propagation approximately every 5-7 days. Once established, the organoid lines expanded rapidly following each passage (**Figure S6B-C, S6F**). Morphologically, Jfb TOs and DOs closely resembled those derived from human and rhesus macaque placentas (**Figure 6C**)^41,42,44^. TOs exhibited a dense, solid morphology, whereas DOs formed cystic structures with a central lumen. H&E staining confirmed these observations, with TOs displaying dense, multilayered 3D structures, some of which contained intra-organoid cavities, consistent with features observed in human TOs^44,45^ (**Figure 6D, 6E**). In contrast, DOs exhibited a hollow, cystic morphology lined by a single layer of epithelial-like cells surrounding the central lumen (**Figure 6F**). While human and Jfb TOs were morphologically similar (**Figure 6G**), they exhibited notable differences in size, with Jfb TOs forming smaller, more compact structures in culture. To quantitatively assess this observation, we developed an unbiased, microscopy-based platform to measure the cross-sectional area of all organoids within individual Matrigel domes. This approach enabled systematic, high-throughput quantification of organoid size across replicate cultures, allowing direct comparison of size distributions between species under standardized culture conditions (**Figure S6D-E**). Across multiple lines, JfbTOs were consistently smaller than human TOs, with significantly reduced cross-sectional areas (**Figure 6H**, **Figure S6D-E**).

**Figure 6.**
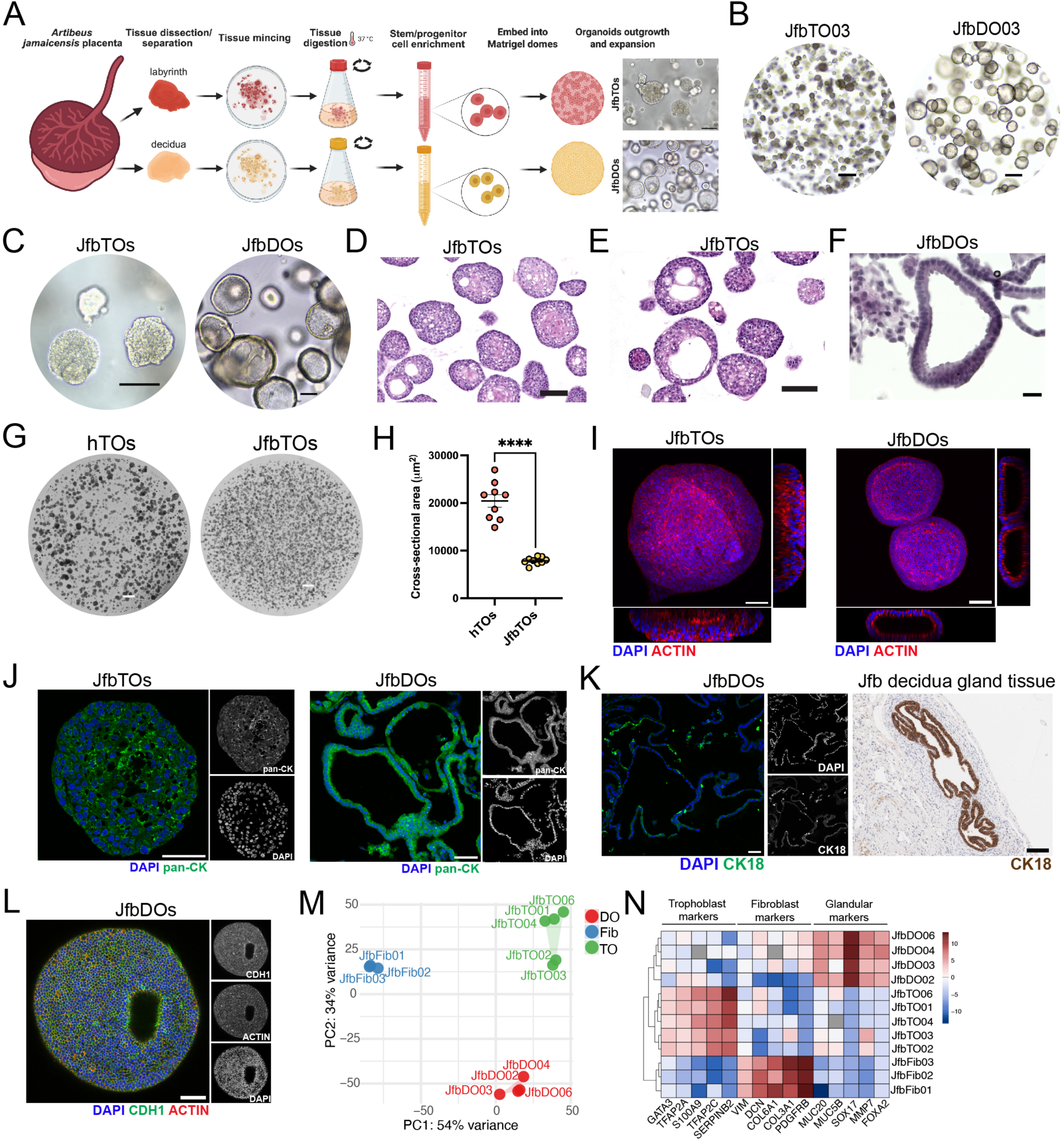
Derivation and Characterization of Trophoblast and Decidual Gland Organoids from the Jamaican Fruit Bat Placenta. **(A)** Schematic of Jfb placental organoid derivation, including dissection of labyrinth (fetal-derived) and decidua (maternal-derived) tissues, enzymatic digestion, and organoid 3D culture in Matrigel. **(B)** Representative brightfield images showing successful derivation of trophoblast organoids (TOs) and decidual organoids (DOs) from Jfb placenta. Scale bars, 500 μm. **(C)** High-magnification brightfield images highlighting the distinct morphologies of established JfbTOs and JfbDOs. Scale bars, 200 μm. **(D–F)** Hematoxylin and eosin (H&E) staining of TOs and DOs, showing dense 3D architecture with internal cavities in TOs (D, E) and a hollow cystic morphology in DOs (F). Scale bars, 100 μm (D, E); 50 μm (F). **(G)** Brightfield comparison of Jfb and human TOs, demonstrating size differences. Scale bars, 500 μm. **(H)** Quantification of organoid cross-sectional areas using high-throughput microscopy, showing that Jfb TOs are significantly smaller than human TOs. Data points represent the average cross-sectional area per field of view (three fields per Matrigel dome). ****p < 0.0001, two-tailed unpaired t-test with Welch’s correction. **(I)** Whole-mount confocal imaging of JfbTOs (left) and JfbDOs (right) stained for actin (red) and nuclei (DAPI, blue), with orthogonal projections below and right. Scale bars, 50 μm. **(J)** Confocal immunofluorescence for pan-cytokeratin (green) confirming epithelial/trophoblast identity of TOs and DOs. Right panels show individual channels in grayscale. Scale bars, 50 μm. **(K)** Confocal immunofluorescence for KRT18 (green) in JfbDOs (left) and immunohistochemistry (IHC) for KRT18 in placental tissue (right), validating glandular epithelial lineage. Nuclei counterstained with DAPI (blue); grayscale individual channels at right. Scale bars, 50 μm (DOs); 100 μm (tissue). **(L)** Confocal immunofluorescence for CDH1 (green) and actin (red) in JfbDOs, further supporting epithelial identity. Nuclei counterstained with DAPI (blue); grayscale individual channels at right. Scale bar, 50 μm. **(M)** Principal component analysis (PCA) of bulk RNA-seq data showing clear separation of TOs (green), DOs (red), and fibroblasts (blue) with tight clustering of biological replicates. **(N)** Heatmap of canonical lineage markers (based on log_2_ RPKM values) confirming trophoblast identity of TOs, glandular epithelial identity of DOs, and mesenchymal identity of fibroblasts.

To verify the cellular identity of Jfb organoids, we next applied immunofluorescence staining for known cytoskeletal and epithelial markers. Actin staining revealed distinct cytoskeletal organization between TOs and DOs, consistent with their divergent morphologies (**Figure 6I**). Staining for pan-CK confirmed that both organoid types were composed of epithelial or trophoblast-derived cells (**Figure 6J**). Given that KRT18 expression was observed in glandular epithelial structures within the decidua of Jfb placental tissue by immunohistochemistry (**Figure 6K**, right), we further confirmed that DOs were KRT18-positive by immunofluorescence (**Figure 6K**, left). In addition, DOs expressed the epithelial adhesion marker CDH1, further supporting their identity as glandular epithelial organoids (**Figure 6L**).

To further confirm the cellular identity and purity of Jfb organoids, we performed bulk RNA sequencing on TOs derived from all five placentas and matched DOs from four placentas. In parallel, we generated transcriptomes from primary placental fibroblasts isolated at the time of organoid derivation from three placentas. This approach allowed us to assess the specificity of each organoid type and to rule out cross-contamination with fibroblasts or other epithelial populations. Principal component analysis (PCA) revealed clear separation between TOs, DOs, and fibroblasts, with tight clustering of biological replicates, indicating both cellular distinctiveness and reproducibility across independent preparations (**Figure 6M**). Hierarchical clustering based on canonical lineage markers further supported these findings: TOs were enriched for trophoblast-associated genes including *GATA3, TFAP2A*, and *TFAP2C*; DOs expressed markers of glandular epithelium such as *MUC20, MUC5B*, and *SOX17*; and fibroblasts showed selective expression of mesenchymal genes including *VIM, DCN*, and *COL6A1* (**Figure 6N**).

To complement these analyses, we performed differential gene expression analysis using DESeq2, which further confirmed the transcriptional distinctiveness of each population (**Figure S7A**). Volcano plots and heatmaps of the top differentially expressed genes highlighted robust separation of TOs, DOs, and fibroblasts based on their lineage-specific transcriptional programs (**Figure S7A, S7B**). As an orthogonal validation, we also performed single-sample gene set enrichment analysis (ssGSEA) using curated marker sets for trophoblast, glandular epithelial, and adventitial fibroblast populations. Signature scores aligned with the expected lineage identities, with TOs exhibiting strong trophoblast scores, DOs for glandular score, and fibroblasts selectively expressing the adventitial fibroblast signature (**Figure S7C**). Together, these results confirm the molecular identity of each organoid type, demonstrate the absence of contaminating cell populations, and validate the fidelity of our organoid derivation protocols.

### Lineage Differentiation and Attenuated Antiviral Responses in Bat Trophoblast Organoids

To define the cellular composition and differentiation potential of Jfb TOs, we performed snRNA-seq, revealing 11 distinct trophoblast clusters (**Figure S8A-S8C**). All clusters expressed canonical markers (*GATA3*, *TFAP2A/C*, *IRF6*, *KRT7*), confirming their trophoblast identity, which was consistent with immunostaining for GATA3 and KRT7 (**Figure 7A, 7B**). To assign cellular identities to these clusters, we used a multi-step strategy integrating probabilistic modeling, trajectory inference, and marker analysis. First, we applied Seurat’s transfer anchor framework to project *in vivo* trophoblast annotations onto TOs, revealing populations corresponding to proliferative progenitors (TB-p), mononuclear trophoblasts (mTB), and syncytiotrophoblasts (sTB) (**Figure S8D**). Next, we used Slingshot to infer lineage relationships, designating the most proliferative cluster (MKI67⁺, Cluster 1, **Figure S8E**) as the root. This analysis revealed four differentiation trajectories terminating in clusters with distinct gene expression profiles (**Figure S8F**). For each lineage path, we extracted the 10% most terminal cells and assessed their gene expression profiles to assign cell type identities (**Figure S8G**). This revealed five distinct endpoints corresponding to two syncytiotrophoblast (sTB) populations, two invasive trophoblast (iTB) populations, and one mononuclear trophoblast (mTB) population (**Figure 7C**). These annotations were based on transcriptional similarity to *in vivo* trophoblast subtypes and expression of defining markers such as *HOPX* and *KRT18* (sTB), *SLC7A11* and *MACF1* (iTB), and *PRKN* and *ATP6V0A1* (mTB) (**Figure S8G**). Intermediate states included three TB-p clusters and three transitioning trophoblasts (tTB), positioned along differentiation trajectories (**Figure 7C**, **S8F**). These lineage assignments were supported by expression of canonical genes, including *TEAD4* and *MSX2* (progenitors), and *KRT18*, *MAL2*, and *HOPX* (sTB), with KRT18⁺ cells localizing to intra-organoid cavities reminiscent of syncytial domains in human TOs (**Figure 7D**, **S8H**). Gene Ontology analysis further supported distinct biological programs across lineages: TB-p clusters were enriched for mitotic progression and chromatin regulation; tTBs for epithelial morphogenesis and cytoskeletal remodeling; iTBs for matrix interaction and vesicle trafficking; and sTBs for lipid metabolism and hormone secretion (**Figure S8I**, **S8K**). Notably, sTB2 expressed autophagy-related genes (*ATG9B*, *MAP1LC3A*), suggesting functional maturation (**Figure S8I**). Together, these data show that Jfb TOs recapitulate key trophoblast differentiation pathways and support reconstruction of lineage hierarchies *in vitro*.

**Figure 7.**
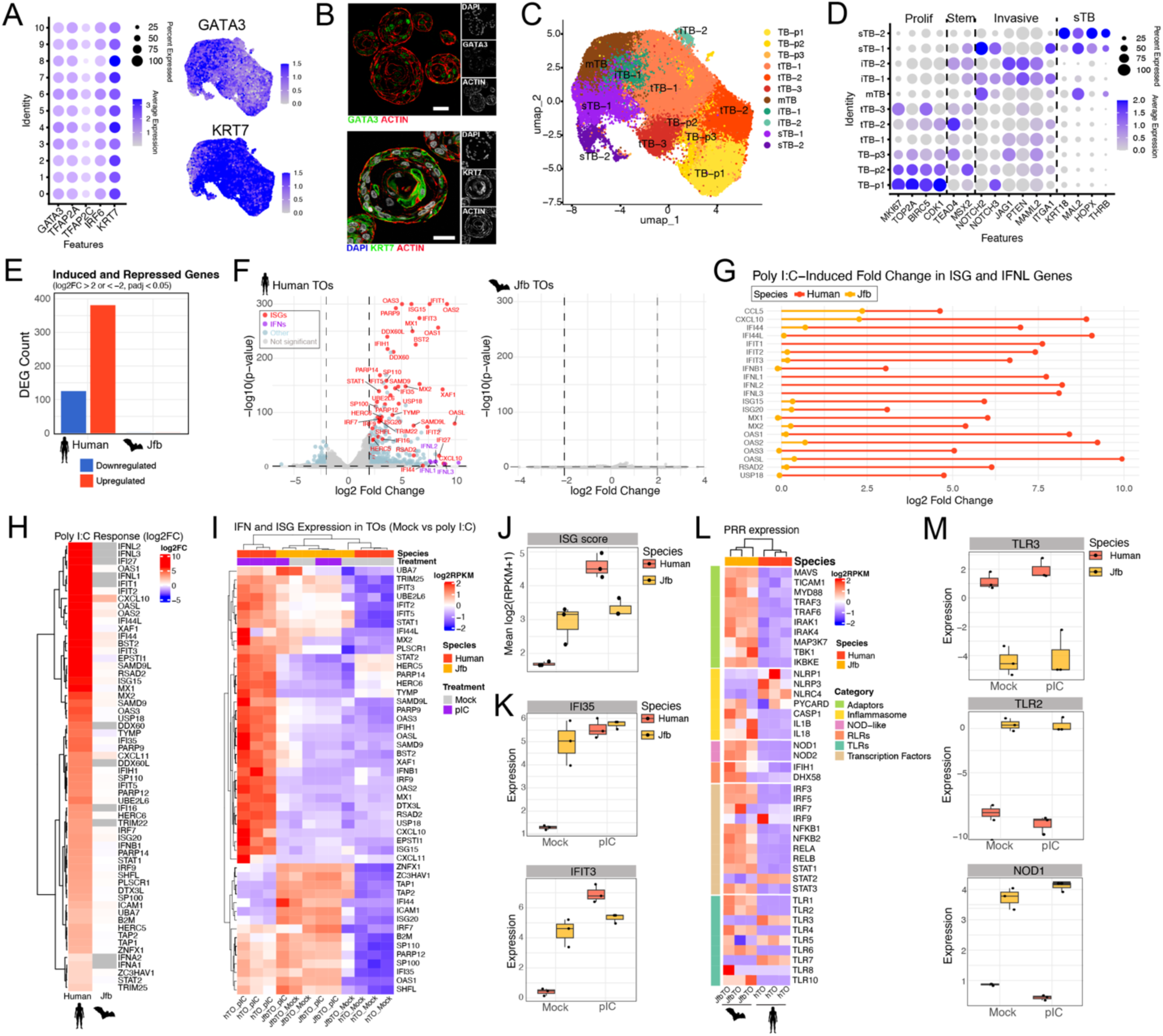
Bat Trophoblast Organoids Recapitulate Placental Lineages and Exhibit Attenuated Antiviral Signaling Compared to Human. **(A)** Left, DotPlot of canonical trophoblast marker expression across TO-derived clusters. Right, FeaturePlots of *GATA3* (top) and *KRT7* (bottom) expression in TOs. Scales at right. **(B)** Confocal immunofluorescence staining for GATA3 (top, green) and KRT7(bottom, green) with actin (red), confirming widespread trophoblast marker expression throughout organoids. Nuclei are counterstained with DAPI (blue); right panels show individual grayscale channels. Scale bar, 50 μm. **(C)** UMAP showing final cell type annotations based on trajectory and root cluster analysis, including mononuclear trophoblasts (mTB), invasive trophoblasts (iTB1, iTB2), syncytiotrophoblasts (sTB1, sTB2), proliferative progenitors (TB-p1–3), and transitional populations (tTB1–3). Key at right. **(D)** Dot plots of canonical trophoblast markers (e.g., *TEAD4*, *MSX2*, *KRT18*, *MAL2*, *HOPX*), validating cell type assignments. Scale at right. **(E)** Bar plot showing the number of differentially expressed genes (DEGs) following poly I:C treatment in human and Jfb TOs, using DESeq2 (log2FC > 1, adjusted *p* < 0.05). Upregulated genes are shown in red and downregulated are shown in blue. **(F)** Volcano plots of DEGs in human (left) and bat (right) TOs after poly I:C exposure. Significantly upregulated interferon-stimulated genes (ISGs) are labeled in red in human TOs, while Jfb TOs show minimal transcriptional changes. Non-ISG DEGs are shown in light blue and non-significant genes are shown in grey. **(G)** Lollipop plot showing poly I:C–induced log2 fold changes in selected interferon-stimulated genes (ISGs) and interferon lambda (*IFNL*) genes in human (red) and Jfb (yellow) TOs. The plot highlights consistently higher induction of antiviral genes in human cells compared to bat cells following stimulation. **(H)** Heatmap of top poly I:C–responsive genes (ranked by log2FC) showing markedly higher induction in human compared to Jfb TOs. Red indicates high indiction and grey/white indicates no or low induction. Scale at right. **(I)** Heatmap showing scaled average expression (Z-score) of ISGs and IFN genes expressed in mock and poly I:C-treated TOs, grouped by species. Bat TOs exhibit limited upregulation following stimulation. Scale at right. **(J)** Boxplot comparing ISG scores in human and Jfb TOs under mock and poly I:C-treated conditions. Jfb TOs show elevated ISG expression at baseline (mock), whereas human TOs exhibit a greater dynamic induction following poly I:C treatment. **(K)** Boxplots showing expression of representative ISGs (*IRF35*, *IFIT3*) across species and conditions. **(L)** Heatmap showing scaled average expression (Z-score) of key innate immune signaling genes including Toll-like receptors (TLRs), RIG-I–like receptors (RLRs), NOD-like receptors (NLRs), adaptor proteins, inflammasome components, and downstream transcription factors across mock-treated Jfb and human TOs. Genes are grouped by functional category, with color-coded annotations at right. Expression values were derived from gene reads. **(M)** Boxplots comparing baseline and induced expression of select PRRs (*TLR3*, *TLR2*, *NOD1*), further supporting attenuated innate immune sensing in bat TOs.

We previously used human TOs to model antiviral signaling at the maternal-fetal interface^41^. To directly compare antiviral responses across species, we performed bulk RNA-seq on human and Jfb TOs following poly I:C stimulation. As expected, poly I:C treatment induced a robust transcriptional response in human TOs, with hundreds of differentially expressed genes (DEGs), including classical interferon-stimulated genes (ISGs) and type III interferons (IFNLs) (**Figure 7E–F**, **Table S10**). In contrast, Jfb TOs exhibited a strikingly muted response, with no genes meeting thresholds for differential expression (**Table S11**). This species-specific difference was evident in both the magnitude and breadth of ISG induction: antiviral genes such as *IFI44L*, *IFIT1*, *ISG15*, *MX1*, and *RSAD2* were strongly upregulated in human TOs but showed little to no induction in bat TOs (**Figure 7G–H**). Consistent with our prior findings, *IFNL* genes represented the predominant interferons induced in human TOs (**Figure 7G**). To assess antiviral signaling baseline, we calculated an ISG score based on the average expression of 30 canonical ISGs. This score revealed significantly higher basal ISG activity in Jfb TOs under mock conditions compared to human TOs, which instead showed a pronounced increase following stimulation (**Figure 7J**). This elevated ISG baseline in bat cells was also evident in expression of individual ISGs, including *IFI35* and *IFIT3* which were consistently higher in mock-treated Jfb TOs than in human counterparts (**Figure 7K**).

To explore potential upstream mechanisms, we examined expression of pattern recognition receptors (PRRs) involved in viral sensing. Human TOs expressed multiple RNA-sensing PRRs, including *TLR3*, *TLR7*, and *IFIH1*, whereas Jfb TOs exhibited markedly reduced or undetectable expression of these receptors (**Figure 7L–M**). In contrast, bat TOs retained expression of certain bacterial PRRs such as *TLR2* and *NOD1*. To evaluate whether these features of TOs are conserved i*n vivo*, we examined expression of PRRs across trophoblast subtypes in the Jfb placenta using snRNA-seq. Consistent with TOs, multiple trophoblast populations, particularly sTB populations, exhibited downregulation of viral sensors such as *TLR3* and *IFIH1*, whereas other trophoblast subsets retained higher PRR expression (**Figure S8K**, **S8L**). In contrast, invasive trophoblasts (iTBs), which are not enriched in TO cultures, exhibited very high expression of nearly all TLRs and cytosolic viral sensors, as well as downstream signaling components, suggesting a distinct immunological profile. Together, these data support a species-specific strategy in bats in which some trophoblasts maintain elevated basal ISG activity but restrict PRR expression and inducibility in differentiated interface cells, potentially limiting inflammation while preserving antiviral vigilance.

## Discussion

Pregnancy in bats occurs under extreme physiological conditions, likely driving unique adaptations in placental structure, function, and immune regulation. Here, we define the cellular landscape of the Jamaican fruit bat placenta using snRNA-seq, revealing diverse trophoblast subtypes, including proliferative, invasive, and syncytial populations, as well as stromal and immune cells with bat-specific transcriptional programs. Distinct macrophage subsets expressed pregnancy-associated regulatory molecules, suggesting specialized roles in placental development and immune tolerance. To further explore trophoblast lineages, we derived organoids from labyrinth and decidual tissues that recapitulate key *in vivo* states and provide experimental access to otherwise inaccessible stages. To facilitate broad exploration of these datasets, we developed an interactive ShinyApp that enables users to visualize gene expression, cell-type annotations, and differential expression results across tissue and organoid samples (available at https://coynelab.shinyapps.io/bat-placenta-explorer/).

The placenta is a diverse organ that varies across species. Bats, the second largest order of mammals with over 1,400 species, exhibit remarkable placental diversity, including a range of structural types and reproductive strategies^46^. Within Chiroptera, placental architecture spans hemochorial, endotheliochorial, and occasionally epitheliochorial forms, with variations in trophoblast invasiveness, placental morphology, and maternal tissue remodeling^3,14^. Previous studies have documented variation in blastocyst implantation, placental positioning, and placental zones^15^, suggesting substantial differences in cellular composition and transcriptional programs among bat lineages. While our work focused on the Jfb, a member of the microbat clade, it provides the first high-resolution molecular and cellular analysis of the bat placenta. Although this study does not capture the full diversity of placental forms across Chiroptera, it establishes a critical foundation for understanding the cellular architecture and gene expression programs that may be shared or specialized across bat lineages. Notably, we identified a bat-specific expansion of pregnancy-specific glycoproteins (PSGs), a subclass of the CEACAM family, expressed in both trophoblast and macrophage populations, highlighting novel placental adaptations that had not been previously characterized in bats. Jfb are microbats, which differ from megabats in size, ecology, and reproductive physiology^15^, factors likely to influence placental structure and function. Future studies extending single-cell/nuclei and transcriptomic approaches across a broader range of species will be essential to map the evolutionary trajectories of bat placentation and uncover the diversity of placental strategies in this highly unique and diverse mammalian order.

Comparative analysis of placental cell types and transcriptional profiles across bat, human, and mouse revealed marked species-specific differences. Trophoblast populations exhibited limited transcriptional overlap, despite the presence of both invasive and syncytial subtypes across species. While the bat placenta shares structural similarities with the labyrinthine mouse placenta, it also displays features characteristic of the human placenta, including a distinct invasive trophoblast population expressing integrins, adhesion molecules, and immunomodulatory factors. These similarities may reflect convergent evolution, as bats like the Jfb exhibit prolonged gestation periods more typical of larger mammals, necessitating prolonged placental support and maternal adaptation. In humans, EVTs are central to implantation, maternal vascular remodeling, and immune regulation^47^. Our data suggest that similar processes are active in the bat placenta, likely mediated by the invasive trophoblast populations we identified. Interestingly, while individual trophoblast clusters displayed distinct transcriptional profiles, differential expression analyses revealed a shared core of genes enriched across all four bat trophoblast lineages. These findings support a model in which bat trophoblasts engage a coordinated molecular program, potentially enabling functional redundancy, resilience, or shared roles in nutrient exchange, stress adaptation, and immune modulation. Pathway analysis of these shared DEGs highlighted bat-specific upregulation of TORC1 signaling, amino acid starvation responses, and cold-stress pathways, features that may reflect adaptation to energy limitation, temperature fluctuation, or oxidative stress during gestation.

In addition to trophoblasts, stromal fibroblasts exhibited species-specific adaptations that may contribute to the unique physiological demands of bat pregnancy. A Jfb-enriched fibroblast population displayed a transcriptional profile consistent with adventitial fibroblasts, a specialized subset of perivascular stromal cells involved in structural support, extracellular matrix remodeling, mechanosensory signaling, and immune regulation. These cells are typically positioned at the interface of vasculature and surrounding tissue, where they integrate mechanical and inflammatory cues to maintain vascular tone and immune homeostasis. In the bat placenta, adventitial fibroblasts may play an expanded role, functioning not only as structural elements but also as local sensors of infection or tissue stress. Notably, they expressed high levels of PRRs and adaptor molecules (**Figure S8K**), consistent with a role in innate immune surveillance, as well as genes involved in neuronal signaling pathways, suggesting a hybrid identity not observed in other mammalian placentas. This dual transcriptional program may enable them to relay environmental and immunological signals, such as oxygen tension, pathogen exposure, or inflammatory mediators, to neighboring trophoblast and endothelial populations. In doing so, these cells could act as integrative hubs for placental adaptation, coordinating tissue remodeling, vascular responses, and immune signaling under dynamic physiological conditions. Their emergence in the bat placenta highlights the possibility that bats have evolved fibroblast subtypes with specialized immunoregulatory functions to support pregnancy in the face of heightened pathogen exposure and metabolic stress.

The immune landscape of the bat placenta also revealed unexpected adaptations that may underlie the ability of bats to sustain pregnancy despite persistent immune and environmental stress. Macrophages in the Jfb placenta exhibited transcriptional features of tissue-resident, M2-like cells, resembling human Hofbauer cells known for roles in angiogenesis, remodeling, and immune tolerance. In Jfb, these cells were localized at the maternal-fetal interface and expressed markers associated with anti-inflammatory and regulatory functions. Remarkably, these placental macrophages expressed a PZP-like gene and a PSG, which are traditionally restricted to trophoblast or maternal decidual cells in other mammals^29^. This represents a significant departure from established paradigms of placental immune regulation. In humans, both PZP and PSGs promote maternal immune tolerance by modulating cytokine responses, supporting Treg differentiation, and dampening inflammatory signaling^48,49^. PZP has also been shown to bind and stabilize misfolded proteins, including amyloid beta peptides linked to preeclampsia and neurodegenerative disease^35^. This chaperone-like activity may be particularly advantageous in bats, which experience repeated cycles of hyperthermia and oxidative stress during flight, conditions known to promote protein misfolding. The expression of both a PZP-like gene and PSGs in bat macrophages suggests that these immune cells may partially recapitulate trophoblast-like signaling functions. This convergence of trophoblast and immune cell gene expression may reflect an evolved strategy to reinforce immune tolerance in the face of continuous microbial exposure, oxidative stress, and metabolic demands. In parallel, we identified a T-cell population exhibiting transcriptional features of naive, effector, and regulatory states, pointing to a dynamic and adaptable T-cell compartment. In mice and humans, NK cells are abundant at the maternal-fetal interface and are essential for regulating trophoblast invasion, vascular remodeling, and maternal immune tolerance of the placenta^50^. We were unable to identify any NK cells by snRNAseq in the bat placenta. This could be an artifact of gestational time as NK cells are most abundant in other species early in gestation but may also indicate a notable immunological difference in the bat placenta^51,52^. Together, these findings reveal a uniquely complex immune niche in the bat placenta, where immune cells appear to adopt hybrid roles that enhance both defense and tolerance, hallmarks of reproductive success in a physiologically extreme context.

This work establishes scalable *in vitro* models of the Jfb maternal-fetal interface, enabling functional analysis of species-specific transcriptional programs that are difficult to access *in vivo*. We derived and fully characterized TOs and DOs from primary tissue histological and immunofluorescence staining, bulk RNA-seq, and snRNA-seq. Single-nucleus profiling revealed distinct trophoblast subpopulations within TOs that recapitulate key *in vivo* features and states, including intermediate and differentiated lineages. These findings demonstrate that Jfb TOs retain the developmental plasticity of their tissue of origin and offer a tractable system for modeling trophoblast biology. Unlike cell lines or explants, organoids provide a self-renewing platform to study dynamic and transitional trophoblast populations that are rarely captured from intact tissue. To date, molecular markers for bat trophoblasts have been poorly defined; our transcriptomic profiling addresses this gap, offering a resource for identifying lineage markers and regulatory pathways. Limited genome annotation remains a challenge in bat species, but homology-based approaches, such as BLAST, ortholog mapping, and de novo transcriptome assembly, will be key to resolving conserved and bat-specific placental genes. Together, these models and datasets establish a foundation for mechanistic studies of bat placentation and provide a platform to investigate the evolution of placental function under extreme physiological conditions.

Jfb trophoblast organoids exhibited blunted transcriptional responses to poly I:C despite high baseline ISG expression, consistent with tonic interferon signaling observed in other bat cells^9,11^. This combination of constitutive priming and low inducibility may reflect a bat-specific antiviral strategy that limits inflammation while preserving defense, particularly beneficial at the maternal-fetal interface, where immune restraint is critical. Compared to human TOs, which robustly express viral sensors such as *TLR3*, *TLR7*, and *IFIH1*, Jfb TOs selectively expressed bacterial PRRs like TLR2 and NOD1 while lacking many key viral sensors, suggesting an evolutionarily rewired sensing program. Single-nucleus RNA-seq of the intact bat placenta revealed this tuning is spatially and functionally compartmentalized: the syncytiotrophoblast, which directly contact maternal blood, downregulated RNA sensors, while invasive trophoblasts, which are at low levels in TOs but abundant *in vivo*, expressed high levels of TLRs and cytosolic sensors, consistent with a sentinel role at tissue boundaries. Notably, bats possess a distinct population of adventitial fibroblasts that express high levels of PRRs and adaptor proteins (**Figure S8K**), a stromal cell type not observed in the human or mouse placenta. These findings suggest that bats redistribute antiviral sensing to stromal and invasive trophoblast compartments, allowing surface-exposed trophoblasts to remain immunologically restrained. This lineage- and species-specific architecture may represent a divergent evolutionary solution to balancing immune vigilance with tolerance in pregnancy.

Together, our work shows that the bat placenta is comprised of a complex and unique cellular landscape, shaped by transcriptional programs that diverge from well-characterized mammalian models. The combination of single-nucleus profiling and organoid-based systems uncovers both conserved and bat-specific features of trophoblast differentiation, including unique invasive subtypes, and immune and stromal compartments with distinct gene expression signatures. Our organoid models preserve transitional cell states and permit functional manipulation, allowing access to regulatory circuits that are difficult to resolve in intact tissue. These findings suggest that bats achieve successful gestation through placental strategies that balance developmental flexibility with immune restraint. More broadly, this work establishes a foundation for investigating how evolution repurposes cellular states and signaling networks to support reproduction in physiologically extreme environments.

## Acknowledgements

We thank the Molecular Genomics Core at the Duke Molecular Physiology Institute (Duke University School of Medicine) for assistance with sequencing, and Madeline Keenen (Duke University) for support in generating the Shiny app. We are also grateful to Kaila Cooley (Duke University) for technical assistance. Some schematics were created with BioRender.com. Animal silhouettes used in figures were obtained from PhyloPic.org under public domain or Creative Commons licenses. Molecular graphics and analyses were performed using UCSF ChimeraX, developed by the Resource for Biocomputing, Visualization, and Informatics at the University of California, San Francisco, with support from NIH grant R01-GM129325 and the Office of Cyber Infrastructure and Computational Biology, NIAID. This work was supported by The Duke Endowment through the Duke Science and Technology Initiative (CBC) and by NIAID grant R24-AI165424 (TS).

## Materials and Methods

### Placental tissue collection and processing

Jamaican fruit bats were housed at Colorado State University in accordance with IACUC protocols and standards. Bats in this colony are housed in a free-flight vivarium and provided fresh fruit, protein and vitamin supplements, and water each day. The colony rooms are cleaned biweekly. Visibly pregnant bats were euthanized, and the entire fetal-placental unit was harvested and photographed. Fetal and placental measurements were obtained using ImageJ. Placental tissues were dissected and divided into four sections and immediately drop-fixed in 10% neutral-buffered formalin for histological analyses, flash-frozen for snRNAseq, or stored in Dulbecco’s Modified Eagle Medium (DMEM) on ice for organoid derivation. C57BL/6J mice (Jackson Labs #000664) were housed at Duke University in accordance with IACUC protocols and standards. Male and female mice were co-housed for 18 days, after which visibly pregnant females were euthanized. Placentas were dissected and drop-fixed in 10% neutral-buffered formalin for histological analyses. A table containing the details of tissue codes used in experiments is provided (**Table S12**).

### Histology of Jfb and mouse placentas

All staining procedures were performed at HistoWiz, Inc, using the Leica Bond RX automated stainer (Leica Microsystems) and a fully automated workflow. Following fixing in 10% neutral-buffered formalin for 48 hours, bat and mouse placentas were shipped to Histowiz in 70% ethanol, embedded in paraffin, and sectioned. Sections were stained with Hemytoxylin & Eosin, Periodic acid–Schiff (PAS) or with antibodies against Pan-Cytokeratin (abcam, ab308262), Vimentin (abcam, ab92547), KRT18 (CK18) (abcam, ab668), CD45 (abcam, ab10558), CD68 (abcam, ab125212), CD3 (abcam, ab16669), F4/80 (Invitrogen, 14-4801-82), Ki67 (abcam, ab155800), or SOX9 (abcam, ab185230). Full-resolution histological images of all stained sections and placental samples are available via HistoWiz using the following links: <u>H&E</u>, <u>PAS</u>, <u>pan-KRT and KRT18,</u> <u>Vimentin</u>, <u>immune cell markers</u>, <u>Ki67</u>, <u>NCAM1</u>, and <u>SOX9</u>.

### Derivation and culture of trophoblast organoids (TOs) from Jamaican fruit bat placental samples

Bat placental tissues were pre-washed and carefully dissected into fetal placental tissue and maternal-derived decidua for isolation. Labyrinth trophoblast stem/progenitor cells were isolated similar to previous protocols^41^. Briefly, collected labyrinth tissue was cut into small pieces and extensively washed, then sequentially digested with 0.2% trypsin-250 (Alfa Aesar, J63993-09)/0.02% EDTA (Sigma-Aldrich E9884-100G) and 1.0 mg/mL collagenase V (Sigma-Aldrich, C9263-100MG) in small glass containers with stir bars inside placed in a shaking 37°C water bath (Grant Instruments, LSB12US) at 100 rpm. Following collagenase Ⅴ digestion, tissues were manually disrupted by forcefully pipetting up and down about 10 times with a 10 mL serological pipette. Solutions from the two sequential digestions were pooled and filtered through a triple-layered gauze, and the flow-through was collected for downstream processing. The collected flow-through were pooled and centrifuged at 600g for 6 minutes. The pellet was resuspended with 1 × RBC lysis buffer (Invitrogen, 00433357) for 5 min at room temperature. Pelleted cells were washed once with Advanced DMEM/F12 medium (Life Technologies, 12634-010) and finally resuspended in appropriate volume of ice-cold growth-factor-reduced Matrigel (Corning 356231). Matrigel “domes” (one 40ul dome/well) were plated into 24-well tissue culture plates (Corning 3526), placed in a 37°C incubator to pre-polymerize for approximately 3 minutes, turned upside down to ensure equal distribution of the isolated cells in domes for another 10 minutes, then carefully overlaid with 500 µL of prewarmed full growth media. Jfb TOs were derived and grown in term trophoblast organoid medium (tTOM) comprised of Advanced DMEM/F12 (Life Technologies, 12634-010) supplemented with 1X B27 (Life Technologies, 17504-044), 1C N2 (Life Technologies, 17502-048), 10% FBS (vol/vol, Cytiva HyClone, SH30070.03), 2 mM GlutaMAX^TM^ supplement (Life Technologies, 35050-061), 100 µg/mL Primocin (InvivoGen, ant-pm-1), 1.25 mM N-Acetyl-L-cysteine (Sigma, A9165), 500 nM A83-01 (Tocris, 2939), 1.5 µM CHIR99021 (Tocris, 4423), 50 ng/mL recombinant human EGF (Gibco, PHG0314), 80 ng/mL recombinant human R-spondin 1 (R & D systems, 4645-RS-100), 100 ng/mL recombinant human FGF2 (Peprotech, 100-18C), 50 ng/mL recombinant human HGF (Peprotech, 100-39), 10mM nicotinamide (Sigma, N0636-100G), 5 µM Y-27632 (Sigma, Y0503-1MG), and 2.5 µM prostaglandin E2 (PGE2, R & D systems, 22-961-0). To passage, JfbTOs were digested using prewarmed TrypLE Express (Gibco, 12605-028) for 8 min in a 37°C shaking water bath at around 190 rpm followed by manual disruption. Disassociated JfbTOs were centrifuged and resuspended with fresh ice-cold growth-factor-reduced Matrigel. To cryopreserve established JfbTO lines, media was removed and Matrigel domes were scraped off and resuspended with CryoStor CS10 stem cell freezing medium (STEMCell Technologies, 07930) and transferred to −80 ℃ for several hours before were deposited into liquid nitrogen tank for long-term storage. Human TOs were cultured as described previously^41^. A table containing the details of organoid codes used in experiments is provided (**Table S12**).

### Derivation and culture of decidua gland organoids (DOs) from Jamaican fruit bat decidua samples

Dissected decidua tissues were minced into small pieces and washed extensively in wash media (RPMI −1640 with 1x Pen/Strep) prior to being digested in prewarmed dissociation media (1.25 U/mL Dispase II (Sigma-Aldrich, D4693)/0.4mg/mL collagenase V (Sigma-Aldrich, C-9263)) in a 37 ℃ shaking water bath at approximate 100 rpm until decidua glands were observed dominating in the remaining tissues under the microscope (after approximately 20 minutes initial incubation, checking every 5 minutes until tissues were digested well as needed). Following digestion, an equal amount of wash media was added and the remaining tissues were forcefully pipetted approximately 10 times with a 10 mL serological pipette for further dissociation. The released decidual glands were collected by filtration with a 100um-strainer (VWR, 732-2759). The collected glands were pelleted by centrifugation at 600g for 6 minutes and treated with 1 × RBC lysis buffer (Invitrogen, 004333). Following washing with Advanced DMEM/F12 medium, dissociated glands were resuspend in ice-cold growth-factor-reduced Matrigel (Corning 356231), and seeded into Matrigel “domes” in 24-well plates (Corning, 3526), Following polymerization, domes were carefully overlaid with 500 µL prewarmed decidua organoid Expansion Medium (ExM) ^45^. ExM was renewed every 2-3 days. Mature decidua organoids were passaged by mechanical disruption following TrypLE Express (ThermoFisher Scientific, 12605010) digestion every 3-5 days. The cryopreservation of established JfbDOs lines was the same as JfbTOs described above.

### Derivation and culture of primary fibroblast from Jfb placental tissues

Primary fibroblast cultures were established during the initial derivation of JfbTOs. To isolate fibroblasts, the culture media was first aspirated, and cells washed once with 1× DPBS. Then, 1 mL of cell recovery solution (Corning, 354253) was added to each well. Cells were scraped and transferred to microcentrifuge tubes placed on ice to depolymerize the Matrigel, as previously described ^44^. Once the Matrigel had fully dissolved, the samples were centrifuged at 200 × g for 2 minutes to pellet the cells. The pellet was washed once with 1× DPBS, then resuspended in DMEM (Corning, 10017CV) supplemented with 10% FBS (Gibco, A56707), 1% penicillin/streptomycin (Gibco, 15140), 1× HyClone™ non-essential amino acids (Cytiva, SH30238.01), and 1 mM sodium pyruvate (Cytiva, SH30239.01).The entire resuspension from 2–3 JfbTO derivation wells was plated into a single well of a 6-well plate. Plates were incubated for 2 hours at 37°C to allow fibroblasts to rapidly attach. After 2 hours, the plates were gently rocked, and the supernatant was removed to eliminate non-adherent cells. The remaining attached Jfb fibroblasts were maintained in fresh media (as described above).

### Organoid cross-sectional area analysis

Whole-dome brightfield scans were acquired using a Keyence BZ-X810 all-in-one fluorescence microscope equipped with a motorized XY stage. A 24-well plate was mounted onto the stage, and the BZ-X800 Viewer software was used to control image acquisition. Under “Normal” scan mode, “Capture Still Images” was selected. In the “Observation and Capture” window, the imaging mode was set to “Brightfield.” Using the motorized stage controls, the target dome was located. Both Z-stack and Stitching functions were enabled in the “Capture Area Setting” menu. The 2X objective lens (PlanApo, NA 0.10) was selected, and four corner points were defined for stitching. Z-stack limits were set to capture the full depth of the dome, and exposure was adjusted prior to acquisition. Captured image sets were processed using Keyence’s built-in image merging tools to generate stitched, full-focus, high-resolution images of each dome. Organoid cross-sectional area was quantified using Keyence BZ-X800 Analysis Software (v1.1.30.19). From each stitched image, three randomly selected 500 × 500 pixel fields of view per dome were analyzed. A threshold was applied to segment organoids from background, and overlapping organoids were separated using the “Separate Objects” function. Partial organoids at the edges were excluded using “Exclude Objects at Screen Edges,” and debris was removed with the “Remove Objects Less Than” filter (threshold: 200-1500 µm² based on the growth stage of organoids). Organoid counts and cross-sectional areas were calculated per field, and average organoid size was reported per field of view.

### Jfb placental organoid collection, embedding, cryosectioning and immunofluorescence staining

Mature JfbTOs and DOs were released from Matrigel domes as previously described^44^. Organoids were washed with 1× PBS and fixed in 4% paraformaldehyde (PFA) at room temperature (RT) for 2 hours with continuous rotation. Following fixation, organoids were incubated in a 20% (wt/vol) sucrose solution at 4□°C overnight to allow them to sink to the bottom of the tube. The next day, the sucrose solution was removed, and approximately 50□μL of 7.5% gelatin/10% sucrose (wt/vol) embedding medium was carefully added. The mixture of organoids and embedding medium was transferred to mini cryomolds (7 × 7 × 5 mm; Simport Scientific, M475) and incubated at 4□°C for 20 minutes to allow polymerization. Samples were then frozen at –80□°C for at least 6 hours prior to cryosectioning. Frozen organoid blocks were transferred to a cryostat chamber equilibrated to –25□°C for 20 minutes. Blocks were then mounted onto precooled specimen discs and placed on the freezing shelf for further solidification before being inserted into the specimen head for sectioning. Cryosections were cut at 10–15□μm thickness and collected onto microscope slides (Fisherbrand, 1255015). Immunofluorescent staining was performed as previously described ^19,41^. The following primary antibodies were used: KRT18 (1:100; Abcam, ab668), pan-KRT (1:100; Abcam, ab308262), KRT7 (1:200; Abcam, ab181598), GATA3 (CST, 5852), E-cadherin (CDH1, Invitrogen, PA5-85088). Secondary antibodies included Multi-rAb CoraLite Plus 488 Goat anti-mouse (Proteintech, RGAM002) and Multi-rAb CoraLite Plus 594 Goat anti-rabbit (Proteintech, RGAR004). Phalloidin conjugates used were Alexa Fluor 594–phalloidin (Invitrogen, A12381) and Alexa Fluor 647–phalloidin (Invitrogen, A22287). Images were captured using an Olympus Fluoview FV3000 inverted confocal microscope. Image contrast was adjusted using Fiji (v2.14.0/1.54f) or Adobe Photoshop (v24.3.0). In some cases, pseudocoloring was applied for optimal visualization using Fiji or Fluoview software. Image analysis and 3D reconstructions were performed using FIJI.

### Whole-mount immunofluorescence staining of Jfb placental organoids

Mature trophoblast organoids were released and collected for fixation as described above. Instead of embedding for cryosectioning, intact organoids were processed directly for whole-mount immunostaining following permeabilization, as previously described^41,44^. All centrifugation steps were replaced by gravity sedimentation to preserve organoid morphology. To prevent compression during mounting, small dots of Vaseline were applied to the corners of each coverslip. Z-stack fluorescent images were acquired using an Olympus Fluoview FV3000 inverted confocal microscope.

### AlphaFold3 Structural Prediction and Visualization

Predicted protein structures for bat, human, and mouse pregnancy-specific glycoproteins (PSGs) were generated using AlphaFold3.0. Structural overlays were performed using ChimeraX (version 1.9)^53^ to align predicted models based on backbone RMSD minimization. Structures were visualized and colored by species for comparison, and movies were generated within ChimeraX to highlight domain conservation and overall architectural similarity across species.

### RNA extraction and bulk RNAseq of JfbTOs, JfbDOs and Jfb Fibroblasts

Total RNA was extracted from established JfbTOs, JfbDOs, and fibroblasts (Jfb Fibroblasts) at early passage (see **Table S12**) with Sigma GenElute Universal total RNA purification kit (Sigma-Aldrich, RNB100) following manufacturer’s instruction. Purified Total RNA concentration and quality was determined by Thermo scientific Nanodrop One spectrophotomer. All total RNA samples submitted for bulk RNA-seq were further run for QC evaluation for their RQN (RNA quality number, >7 for all samples) prior to library preparation by the Duke Sequencing and Genomic Technologies (SGT) using KAPA HyperPrep kit (Roche). Libraries were sequenced on the NovaSeq X Plus 10B lane. The reads were aligned to the Jfb (*Artibeus jamaicensis*) genome assembly (GCF_021234435.1_CSHL_Jam_final_genomic.fna) using the Rsubread package (v2.10.0) in R version 4.1^54^. Alignment indices were first generated with *buildindex()*, and paired-end reads were aligned using *align()* with four threads. Gene-level quantification was performed using *featureCounts()* (Rsubread) with the associated genome annotation file (GCF_021234435.1, GTF format). Genes were counted based on exon features, and counts were generated for each sample. To facilitate downstream normalization, reads per kilobase per million mapped reads (RPKM) values were calculated for each sample, using gene lengths extracted from the annotation file and the total number of mapped reads per sample. Following quantification, differential expression analysis was performed using the DESeq2 package (v1.34.0) in R^55^. Principal Component Analysis (PCA) was conducted using the *prcomp()* function to visualize major sources of variance between conditions. Heatmaps of log₂-transformed RPKM values were generated with the pheatmap package (v1.0.12) in R^56^. Volcano plots highlighting differentially expressed genes were created using GraphPad Prism (version 9.0). Files associated with bulk RNA-seq studies have been deposited into Sequence Read Archive (PRJNA1251670).

To quantify enrichment of lineage-specific transcriptional programs, we performed single-sample gene set enrichment analysis (ssGSEA) on bulk RNA-seq data from TOs, DOs, and fibroblast cultures. Raw counts were normalized using the DESeq2 package with rlog transformation. Curated gene sets were compiled for adventitial fibroblasts (e.g., *PDGFRA*, *PIEZO2*, *IGF1*), trophoblasts (e.g., *GATA3*, *TFAP2C*, *KRT7*), and glandular epithelial cells (e.g., *PRL*, *LRP2*, *SOX17*). ssGSEA was implemented using the GSVA package, and scores were compared across sample types using one-way ANOVA.

Jfb and human TOs were treated with 10□µg/mL high molecular weight poly I:C (Invivogen, tlrl-pic) or mock-treated for ∼24 hours. Total RNA was extracted as described above. RNA-seq libraries were prepared using the Watchmaker mRNA library prep kit and sequenced on an Illumina NovaSeq X Plus platform (∼70 million 50bp paired-end reads per sample). Reads were aligned to the *Artibeus jamaicensis* or GRCh38 genome using STAR, and gene-level counts were generated with featureCounts. Differential gene expression was analyzed using DESeq2. Technical replicates were grouped by biological replicate (Code) and modeled with a paired design (∼ Code + Treatment). Differentially expressed genes were identified by comparing poly I:C–treated versus mock-treated TOs, with significance defined as adjusted p-value < 0.05.

### Organoid and Tissue Processing for Single-Nuclei RNA Sequencing

To create a single cell suspension of organoids, mature trophoblast organoids were collected at early passage (see **Table S12**) and incubated with TrypLE Express (Invitrogen, 12605036) in a 37 ℃ shaking water bath at approximately 190 rpm for 15 min. Organoids were pelleted by centrifugation at 1250 rpm for 3 minutes and resuspended with 200µL Advanced DMEM/F12 supplemented with 2 mM GlutaMAX supplement, 10 mM HEPES (Gibco, 15630-106), and 1 × Penicillin/Streptomycin (Lonza, 17-602E) (basal media). Then organoids were manually disrupted using a single channel p200 pipette for 3 minutes followed by the addition of 800 µL basal media. Dissociated organoids were pelleted by centrifugation at 1250rpm for 5 minutes and the pellet was resuspended in approximately 250 µL of 1x PBS containing 1% FBS. Sequencing was performed on four organoid lines from unique placental tissues.

For single nuclei RNA-sequencing, flash frozen placentas, with uterus removed, or JfbTOs dissociated as described above were subjected to RNA QC and nuclear isolation using 10x Genomics’ Nuclei Isolation protocol (10x Genomics – Pleasanton, VA) per the manufacturer’s instructions. RNA quality was assessed using an Agilent 4200 TapeStation System. Four placental samples and four organoid samples (see **Table S12**) had an RNA integrity score (RIN) >7 and were used for downstream nuclear isolation. To isolate nuclei, 10mg of minced frozen tissue or a single dome of organoids dissociated into single cells, were resuspended in 0.11x lysis buffer (275uL Lysis Buffer + 0.275uL Reducing Agent B + 2.75uL Surfactant + 2502.23uL 1x chilled PBS) and homogenized using a pestle (approximately 60-70x), prior to incubation on ice for 3-5 minutes. Homogenized samples were then washed twice with nuclei wash and resuspension buffer, followed by centrifugation at 500rcf, 4°C, for 10 minutes. Supernatants were then removed, and nuclei resuspended in appropriate volume of wash and resuspension buffer to a concentration of approximately 1.5 x 10^5^ nuclei/mL, which was determined on a Cellometer Ascend Automated Cell Counter (RevvityWaltham, MA). If necessary, nuclei were filtered through 40µm Flowmi Cell Strainers (Bel-Art – Wayne, NJ) to remove clumps and debris prior to proceeding with the single nuclei assay. Nuclei were visualized at 40x magnification on a LifeTech EVOS FL microscope (Thermo Fisher Scientific – Waltham, MA) to determine quality via nuclear membrane intactness. All samples examined in this study had a nuclear intactness ≥80% and a viability <1% to ensure high quality nuclear isolations. 20,000 single nuclei of each sample were loaded with a Chromium Reagent Kit v4. Tissue libraries were sequenced on a 10B flow cell of the Nova Seq X Plus at a targeted depth of 70,000 reads per nucleus. Cellranger was then used to align reads to the Jamaican Fruit Bat genome (GCF_021234435.1) and create a counts matrix. Files associated with snRNA-seq studies have been deposited into Sequence Read Archive (PRJNA1251203 and PRJNA1251235).

### Single-Nucleus RNA Sequencing Data Processing and Integration

To analyze snRNA-seq data from placental tissue and TOs, we first processed and merged individual datasets separately by sample type. Mitochondrial gene expression was assessed using a curated list of mitochondrial genes (*ND5*, *ND4*, *ND3*, *ND1*, *ND2*, *ND6*, *ND4L*, *CYTB*, *COX2*, *ATP8*, *ATP6*, *COX3*, *COX1*), and mitochondrial and ribosomal RNA content were calculated using the *PercentageFeatureSet()* function. Quality control filtering was tailored to each dataset: for placenta tissue, we retained nuclei with 300–9000 detected genes, fewer than 20,000 UMI counts, ribosomal RNA content below 1.2%, and mitochondrial RNA content below 1.5%. For TOs, dataset-specific thresholds were applied, generally retaining nuclei with 300–9000 genes, fewer than 40,000 UMI counts, ribosomal RNA content below 9%, and mitochondrial RNA content below 2%.

All analyses were conducted in R version 4.3.2 using Seurat v5.0.1, SeuratWrappers v0.3.0 for Harmony integration, and MAST v1.26.0. For figure generation and analysis, ComplexUpset v1.3.3 and ggplot2 v3.5.0 were also used. Sequencing data were aligned and quantified using Cell Ranger v6.1.2 (10x Genomics) against the Jamaican fruit bat genome (annotation release 100), GRCh38 for human, and GRCm39 for mouse. Following quality control, data normalization and dimensionality reduction were performed using the Seurat v5 pipeline. Layers within each dataset were merged using *JoinLayers()* and split by sample identity. We applied *SCTransform()* to normalize gene expression and regress out unwanted technical variation, including gene count, UMI count, and the percentages of mitochondrial and ribosomal RNA. Dimensionality reduction was performed with principal component analysis (PCA) using *RunPCA(),* and the number of components used for downstream analysis was determined using an elbow plot *(ElbowPlot()).* To integrate data across samples while correcting for batch effects, we used Harmony-based integration for placenta tissue (IntegrateLayers() with HarmonyIntegration()) and canonical correlation analysis (CCA) for TOs. In both cases, PCA was replaced with a batch-corrected low-dimensional space for downstream analysis. Clustering was performed using *FindNeighbors()* and *FindClusters()* with a resolution of 0.8 for placenta and 0.2 for TOs. Higher resolution was used for tissue to resolve interstitial and immune subpopulations, while a lower resolution was chosen for organoids to prevent over-clustering of compact epithelial lineages. Batch-corrected UMAP embeddings were generated using *RunUMAP()* to visualize the final integrated structure of each dataset.

Differential gene expression analysis was conducted using *FindAllMarkers()* on the normalized and scaled RNA assay. Marker genes were defined using a minimum log fold change threshold of 0.25 and required expression in at least 25% of nuclei within a cluster (min.pct = 0.25). Marker genes were identified separately for placenta and TO datasets. To address sparsity in gene expression, we applied Adaptively-thresholded Low-Rank Approximation (ALRA) using *RunALRA()* on the SCT assay to impute missing values. The processed and integrated placenta and TO datasets were retained for all downstream analyses.

To assess enrichment of lineage-specific programs in fibroblast populations from the single-nucleus RNA-seq dataset, we computed per-cell signature scores for adventitial fibroblasts and neuronal-associated gene sets. A curated adventitial signature (e.g., *PDGFRA, PIEZO2, IGF1*) and a neuronal-like signature (e.g., *PRICKLE1, MAPK10, GPC4*) were assembled based on published markers and observed gene enrichments in bat-specific fibroblasts (bFib). Signature scores were calculated using Seurat’s *AddModuleScore()* function and aggregated by fibroblast cluster. Score distributions were visualized using boxplots, and statistical comparisons were performed using Wilcoxon rank-sum tests. Full gene lists are provided in **Table S9**.

Differential gene expression analysis was performed using the MAST framework to compare each bat trophoblast cluster (bTB-1 through bTB-4) against human and mouse trophoblasts. Genes were considered significantly upregulated if they met a log₂ fold-change > 2 and adjusted p-value < 0.05. To visualize shared and cluster-specific differentially expressed genes, we generated an intersection plot using the ComplexUpset R package. A binary matrix of gene-cluster associations was constructed, and intersections across clusters were visualized, highlighting the conserved transcriptional program shared among bat trophoblast populations.

Custom code used for data analysis and figure generation is available at CoyneLabDuke GitHub repository.

### Cross-species Single-Nucleus RNA Sequencing Integration and Analysis

To enable integrated analysis across species, we processed snRNA-seq datasets from Jfb placenta, first-trimester human placenta, and mid-gestation (E12) mouse placenta using a standardized workflow. Because gene symbols differ in case formatting across species, we standardized mouse gene names to uppercase to match bat and human gene naming conventions. This was done by extracting the RNA assay count matrix from the mouse Seurat object, converting all gene names to uppercase using toupper(), and removing version suffixes (e.g., ".1", ".2") using regular expressions. To avoid duplicated gene names after case conversion, a custom function was applied to append unique identifiers (e.g., “_DUP1”, “_DUP2”) to duplicated entries. The modified count matrix was then used to generate a new Seurat object with standardized gene names, and UMAP embeddings, cluster identities, and metadata from the original mouse object were transferred to preserve downstream analyses. RNA and ALRA assays were also recreated in the renamed object to ensure consistency across layers.

For integration, shared genes were identified across all three species by computing pairwise intersections of RNA assay gene sets using *LayerData()* and *intersect()* (base R). This resulted in a set of 13,546 genes shared across bat, human, and mouse datasets. Each object was subset to retain only these genes, ensuring compatibility across species. Normalization was performed independently for each dataset using *SCTransform()* v2 in Seurat v5, with regression of covariates including gene count, UMI count, and mitochondrial and ribosomal RNA percentages. Normalized data were stored in new SCT assay layers. The normalized datasets were then merged using *merge()* to form a unified Seurat object. Dimensionality reduction was performed using PCA (*RunPCA()),* and integration was carried out using *IntegrateLayers()* with HarmonyIntegration, using species identity as the batch variable. The Harmony-reduced space was used for neighbor detection (*FindNeighbors()),* clustering (*FindClusters(),* resolution = 0.3), and visualization (*RunUMAP()),* enabling integrated cross-species analysis of conserved and divergent placental cell types.

To preserve species-specific transcriptomic complexity, Harmony and UMAP embeddings from the integrated object were projected back onto the original, full-genome Seurat objects using *Embeddings()* and *CreateDimReducObject()* with species-specific assay settings. Only shared cells between the original and integrated datasets were retained. Harmony embeddings were added to the SCT assay, and UMAP coordinates to the RNA assay, using distinct keys (“harmony_”, “UMAPharmony_”). Each dataset was then independently clustered using Harmony reduction and FindClusters() (resolution = 0.3), allowing for visualization and downstream analysis of integrated cell states while retaining the full gene expression profile of each species.

To annotate predicted genes labeled as "LOC" identifiers in the *Artibeus jamaicensis* genome, we developed a custom R-based pipeline. LOC IDs were queried against the NCBI Gene database using the rentrez R package^57^ to retrieve corresponding official gene names, functional descriptions, and gene types. LOC IDs were queried against the NCBI Gene database using the rentrez R package^57^ to retrieve corresponding official gene names, functional descriptions, and gene types. For each LOC ID, we first retrieved the associated NCBI Gene ID, then extracted the gene name, description, and annotation status (e.g., validated, provisional). The resulting annotated gene list was merged with the original LOC dataset and exported for downstream analysis.

Parts of the analysis code and Shiny app interface were developed with assistance from ChatGPT (OpenAI). Custom code used for data analysis is available at <u>CoyneLabDuke GitHubrepository</u> and an interactive ShinyApp for exploring snRNA-seq and bulk RNA-seq data from this study is available at https://coynelab.shinyapps.io/bat-placenta-explorer/.

## Literature Cited

1. Enders, A.C., and Carter, A.M. (2004). What Can Comparative Studies of Placental Structure Tell Us?—A Review. Placenta 25, S3–S9. 10.1016/j.placenta.2004.01.011.

2. Roberts, R.M., Green, J.A., and Schulz, L.C. (2016). The evolution of the placenta. Reproduction 152, R179–R189. 10.1530/rep-16-0325.

3. Crichton, E.G., and Krutzsch, P.H. (2000). Reproductive Biology of Bats (Academic Press). 10.1016/B978-0-12-195670-7.X5000-0.

4. Willis, C., Brigham, R., and Geiser, F. (2006). Deep, prolonged torpor by pregnant, free-ranging bats. Die Naturwissenschaften 93, 80–83. 10.1007/s00114-005-0063-0.

5. Brook, C.E., and Dobson, A.P. (2015). Bats as ‘special’ reservoirs for emerging zoonotic pathogens. Trends Microbiol 23, 172–180. 10.1016/j.tim.2014.12.004.

6. Schountz, T. (2014). Immunology of bats and their viruses: challenges and opportunities. Viruses 6, 4880–4901. 10.3390/v6124880.

7. Ruiz-Aravena, M., McKee, C., Gamble, A., Lunn, T., Morris, A., Snedden, C.E., Yinda, C.K., Port, J.R., Buchholz, D.W., Yeo, Y.Y., et al. (2022). Ecology, evolution and spillover of coronaviruses from bats. Nat Rev Microbiol 20, 299–314. 10.1038/s41579-021-00652-2.

8. Wong, S., Lau, S., Woo, P., and Yuen, K.Y. (2007). Bats as a continuing source of emerging infections in humans. Rev Med Virol 17, 67–91. 10.1002/rmv.520.

9. Zhou, P., Tachedjian, M., Wynne, J.W., Boyd, V., Cui, J., Smith, I., Cowled, C., Ng, J.H., Mok, L., Michalski, W.P., et al. (2016). Contraction of the type I IFN locus and unusual constitutive expression of IFN-alpha in bats. Proc Natl Acad Sci U S A 113, 2696–2701. 10.1073/pnas.1518240113.

10. Xie, J., Li, Y., Shen, X., Goh, G., Zhu, Y., Cui, J., Wang, L.F., Shi, Z.L., and Zhou, P. (2018). Dampened STING-Dependent Interferon Activation in Bats. Cell Host Microbe 23, 297–301 e294. 10.1016/j.chom.2018.01.006.

11. Ahn, M., Anderson, D.E., Zhang, Q., Tan, C.W., Lim, B.L., Luko, K., Wen, M., Chia, W.N., Mani, S., Wang, L.C., et al. (2019). Dampened NLRP3-mediated inflammation in bats and implications for a special viral reservoir host. Nat Microbiol 4, 789–799. 10.1038/s41564-019-0371-3.

12. Wimsatt, W.A. (1954). The fetal membranes and placentation of the tropical American vampire bat Desmodus rotundus murinus, with notes on the histochemistry of the placenta. Acta Anat (Basel) 21, 285–341. 10.1159/000140938.

13. Carter, A.M., and Enders, A.C. (2004). Comparative aspects of trophoblast development and placentation. Reprod Biol Endocrinol 2, 46. 10.1186/1477-7827-2-46.

14. Badwaik, N., and Rasweiler, I. (2000). Pregnancy. Reproductive biology of bats, 221–293.

15. Carter, A.M., and Mess, A. (2008). Evolution of the placenta and associated reproductive characters in bats. J Exp Zool B Mol Dev Evol 310, 428–449. 10.1002/jez.b.21216.

16. Cukierski, M.A. (1987). Synthesis and transport studies of the intrasyncytial lamina: an unusual placental basement membrane in the little brown bat, Myotis lucifugus. Am J Anat 178, 387–409. 10.1002/aja.1001780409.

17. Ventura, A., Nogueira, M.R., Peracchi, A.L., do Nascimento, A.A., Vieira-Lopes, D.A., and Pinheiro, N.L. (2018). Comparative prenatal development and embryonic staging of neotropical fruit bats (genus Artibeus). Zoologischer Anzeiger 274, 131–146. 10.1016/j.jcz.2018.01.008.

18. Isaac, S.M., Langford, M.B., Simmons, D.G., and Adamson, S.L. (2014). Anatomy of the Mouse Placenta Throughout Gestation. In The guide to investigation of mouse pregnancy, B.A. Croy, A.T. Yamada, F.J. DeMayo, and S.L. Adamson, eds. (Academic Press), pp. 832.

19. Keenen, M.M., Yang, L., Liang, H., Farmer, V.J., Singh, R., Gladfelter, A.S., and Coyne, C.B. (2024). Comparative analysis of the syncytiotrophoblast in placenta tissue and trophoblast organoids using snRNA sequencing. bioRxiv. 10.1101/2024.07.01.601571.

20. Wang, M., Liu, Y., Sun, R., Liu, F., Li, J., Yan, L., Zhang, J., Xie, X., Li, D., Wang, Y., et al. (2024). Single-nucleus multi-omic profiling of human placental syncytiotrophoblasts identifies cellular trajectories during pregnancy. Nat Genet 56, 294–305. 10.1038/s41588-023-01647-w.

21. Vento-Tormo, R., Efremova, M., Botting, R.A., Turco, M.Y., Vento-Tormo, M., Meyer, K.B., Park, J.E., Stephenson, E., Polanski, K., Goncalves, A., et al. (2018). Single-cell reconstruction of the early maternal-fetal interface in humans. Nature 563, 347–353. 10.1038/s41586-018-0698-6.

22. Liu, Y., Fan, X., Wang, R., Lu, X., Dang, Y.L., Wang, H., Lin, H.Y., Zhu, C., Ge, H., Cross, J.C., and Wang, H. (2018). Single-cell RNA-seq reveals the diversity of trophoblast subtypes and patterns of differentiation in the human placenta. Cell Res 28, 819–832. 10.1038/s41422-018-0066-y.

23. Pavlicev, M., Wagner, G.P., Chavan, A.R., Owens, K., Maziarz, J., Dunn-Fletcher, C., Kallapur, S.G., Muglia, L., and Jones, H. (2017). Single-cell transcriptomics of the human placenta: inferring the cell communication network of the maternal-fetal interface. Genome Res 27, 349–361. 10.1101/gr.207597.116.

24. Wang, Q., Li, J., Wang, S., Deng, Q., An, Y., Xing, Y., Dai, X., Li, Z., Ma, Q., Wang, K., et al. (2022). Single-cell transcriptional profiling reveals cellular and molecular divergence in human maternal-fetal interface. Sci Rep 12, 10892. 10.1038/s41598-022-14516-z.

25. Carter, A.M. (2022). Evolution of Placental Hormones: Implications for Animal Models. Front Endocrinol (Lausanne) 13, 891927. 10.3389/fendo.2022.891927.

26. Moore, T., Williams, J.M., Becerra-Rodriguez, M.A., Dunne, M., Kammerer, R., and Dveksler, G. (2022). Pregnancy-specific glycoproteins: evolution, expression, functions and disease associations. Reproduction 163, R11–R23. 10.1530/REP-21-0390.

27. Kromer, B., Finkenzeller, D., Wessels, J., Dveksler, G., Thompson, J., and Zimmermann, W. (1996). Coordinate expression of splice variants of the murine pregnancy-specific glycoprotein (PSG) gene family during placental development. Eur J Biochem 242, 280–287. 10.1111/j.1432-1033.1996.0280r.x.

28. Lei, K.J., Sartwell, A.D., Pan, C.J., and Chou, J.Y. (1992). Cloning and expression of genes encoding human pregnancy-specific glycoproteins. J Biol Chem 267, 16371–16378.

29. Moore, T., and Dveksler, G.S. (2014). Pregnancy-specific glycoproteins: complex gene families regulating maternal-fetal interactions. Int J Dev Biol 58, 273–280. 10.1387/ijdb.130329gd.

30. Rebstock, S., Lucas, K., Weiss, M., Thompson, J., and Zimmermann, W. (1993). Spatiotemporal expression of pregnancy-specific glycoprotein gene rnCGM1 in rat placenta. Dev Dyn 198, 171–181. 10.1002/aja.1001980303.

31. Hallgren, J., Tsirigos, K.D., Pedersen, M.D., Almagro Armenteros, J.J., Marcatili, P., Nielsen, H., Krogh, A., and Winther, O. (2022). DeepTMHMM predicts alpha and beta transmembrane proteins using deep neural networks. bioRxiv, 2022.2004.2008.487609. 10.1101/2022.04.08.487609.

32. Waterhouse, A.M., Procter, J.B., Martin, D.M., Clamp, M., and Barton, G.J. (2009). Jalview Version 2--a multiple sequence alignment editor and analysis workbench. Bioinformatics 25, 1189–1191. 10.1093/bioinformatics/btp033.

33. Tamura, K., Stecher, G., and Kumar, S. (2021). MEGA11: Molecular Evolutionary Genetics Analysis Version 11. Mol Biol Evol 38, 3022–3027. 10.1093/molbev/msab120.

34. Abramson, J., Adler, J., Dunger, J., Evans, R., Green, T., Pritzel, A., Ronneberger, O., Willmore, L., Ballard, A.J., Bambrick, J., et al. (2024). Accurate structure prediction of biomolecular interactions with AlphaFold 3. Nature 630, 493–500. 10.1038/s41586-024-07487-w.

35. Cater, J.H., Kumita, J.R., Zeineddine Abdallah, R., Zhao, G., Bernardo-Gancedo, A., Henry, A., Winata, W., Chi, M., Grenyer, B.S.F., Townsend, M.L., et al. (2019). Human pregnancy zone protein stabilizes misfolded proteins including preeclampsia- and Alzheimer’s-associated amyloid beta peptide. Proc Natl Acad Sci U S A 116, 6101–6110. 10.1073/pnas.1817298116.

36. Ekelund, L., and Laurell, C.B. (1994). The pregnancy zone protein response during gestation: a metabolic challenge. Scand J Clin Lab Invest 54, 623–629. 10.3109/00365519409087542.

37. Lopez-Tello, J., Yong, H.E.J., Sandovici, I., Dowsett, G.K.C., Christoforou, E.R., Salazar-Petres, E., Boyland, R., Napso, T., Yeo, G.S.H., Lam, B.Y.H., et al. (2023). Fetal manipulation of maternal metabolism is a critical function of the imprinted Igf2 gene. Cell Metab 35, 1195–1208 e1196. 10.1016/j.cmet.2023.06.007.

38. Guo, J., Zheng, J., Li, R., Yao, J., Zhang, H., Wang, X., and Zhang, C. (2024). Single-cell transcriptome analysis reveals abnormal angiogenesis and placentation by loss of imprinted glutaminyl-peptide cyclotransferase. Journal of Zhejiang University-SCIENCE B.

39. Buechler, M.B., Pradhan, R.N., Krishnamurty, A.T., Cox, C., Calviello, A.K., Wang, A.W., Yang, Y.A., Tam, L., Caothien, R., Roose-Girma, M., et al. (2021). Cross-tissue organization of the fibroblast lineage. Nature 593, 575–579. 10.1038/s41586-021-03549-5.

40. Pham, T.X., Lee, J., Guan, J., Caporarello, N., Meridew, J.A., Jones, D.L., Tan, Q., Huang, S.K., Tschumperlin, D.J., and Ligresti, G. (2022). Transcriptional analysis of lung fibroblasts identifies PIM1 signaling as a driver of aging-associated persistent fibrosis. JCI Insight 7. 10.1172/jci.insight.153672.

41. Yang, L., Semmes, E.C., Ovies, C., Megli, C., Permar, S., Gilner, J.B., and Coyne, C.B. (2022). Innate immune signaling in trophoblast and decidua organoids defines differential antiviral defenses at the maternal-fetal interface. Elife 11. 10.7554/eLife.79794.

42. Caldwell, A., Yang, L., Scheef, E.A., Kaur, A., and Coyne, C.B. (2024). Comparative analysis of rhesus macaque and human placental organoids highlights evolutionary differences in placentation. bioRxiv, 2024.2010.2011.617873. 10.1101/2024.10.11.617873.

43. McCutcheon, C.R., Caldwell, A., Yang, L., Crisci, E., Pasternak, J.A., and Coyne, C.B. (2024). Defining Cellular Diversity at the Swine Maternal-Fetal Interface Using Spatial Transcriptomics and Organoids. bioRxiv, 2024.2010.2021.619461. 10.1101/2024.10.21.619461.

44. Yang, L., Liang, P., Yang, H., and Coyne, C.B. (2024). Trophoblast organoids with physiological polarity model placental structure and function. J Cell Sci 137. 10.1242/jcs.261528.

45. Turco, M.Y., Gardner, L., Hughes, J., Cindrova-Davies, T., Gomez, M.J., Farrell, L., Hollinshead, M., Marsh, S.G.E., Brosens, J.J., Critchley, H.O., et al. (2017). Long-term, hormone-responsive organoid cultures of human endometrium in a chemically defined medium. Nat Cell Biol 19, 568–577. 10.1038/ncb3516.

46. Rodriguez, F.E., Sandoval, M.T., Alvarez, B.B., and Lombardo, D.M. (2018). Comparative Study of Prenatal Development Between Myotis albescens (Chiroptera: Vespertilionidae) and Eumops patagonicus (Chiroptera: Molossidae): The Chorionic Vesicle and Extraembryonic Membranes Considerations. Anat Rec (Hoboken) 301, 1527–1543. 10.1002/ar.23896.

47. Wei, X.W., Zhang, Y.C., Wu, F., Tian, F.J., and Lin, Y. (2022). The role of extravillous trophoblasts and uterine NK cells in vascular remodeling during pregnancy. Front Immunol 13, 951482. 10.3389/fimmu.2022.951482.

48. Snyder, S.K., Wessner, D.H., Wessells, J.L., Waterhouse, R.M., Wahl, L.M., Zimmermann, W., and Dveksler, G.S. (2001). Pregnancy-specific glycoproteins function as immunomodulators by inducing secretion of IL-10, IL-6 and TGF-beta1 by human monocytes. Am J Reprod Immunol 45, 205–216. 10.1111/j.8755-8920.2001.450403.x.

49. Jones, K., Ballesteros, A., Mentink-Kane, M., Warren, J., Rattila, S., Malech, H., Kang, E., and Dveksler, G. (2016). PSG9 Stimulates Increase in FoxP3+ Regulatory T-Cells through the TGF-beta1 Pathway. PLoS One 11, e0158050. 10.1371/journal.pone.0158050.

50. Moffett, A., and Colucci, F. (2014). Uterine NK cells: active regulators at the maternal-fetal interface. J Clin Invest 124, 1872–1879. 10.1172/JCI68107.

51. Whettlock, E.M., Woon, E.V., Cuff, A.O., Browne, B., Johnson, M.R., and Male, V. (2022). Dynamic Changes in Uterine NK Cell Subset Frequency and Function Over the Menstrual Cycle and Pregnancy. Front Immunol 13, 880438. 10.3389/fimmu.2022.880438.

52. Sojka, D.K., Yang, L., and Yokoyama, W.M. (2019). Uterine Natural Killer Cells. Front Immunol 10, 960. 10.3389/fimmu.2019.00960.

53. Meng, E.C., Goddard, T.D., Pettersen, E.F., Couch, G.S., Pearson, Z.J., Morris, J.H., and Ferrin, T.E. (2023). UCSF ChimeraX: Tools for structure building and analysis. Protein Sci 32, e4792. 10.1002/pro.4792.

54. Liao, Y., Smyth, G.K., and Shi, W. (2019). The R package Rsubread is easier, faster, cheaper and better for alignment and quantification of RNA sequencing reads. Nucleic Acids Res 47, e47. 10.1093/nar/gkz114.

55. Love, M.I., Huber, W., and Anders, S. (2014). Moderated estimation of fold change and dispersion for RNA-seq data with DESeq2. Genome Biol 15, 550. 10.1186/s13059-014-0550-8.

56. Kolde, R. (2018). pheatmap: Pretty Heatmaps. https://github.com/raivokolde/pheatmap.

57. Winter, D.J. (2017). rentrez: an R package for the NCBI eUtils API. The R Journal 9, 520–526.

