## Supplemental Figures and Legends for "Cellular and Immune Adaptations at the Maternal–Fetal Interface in Bats"

**Figure S1**

A

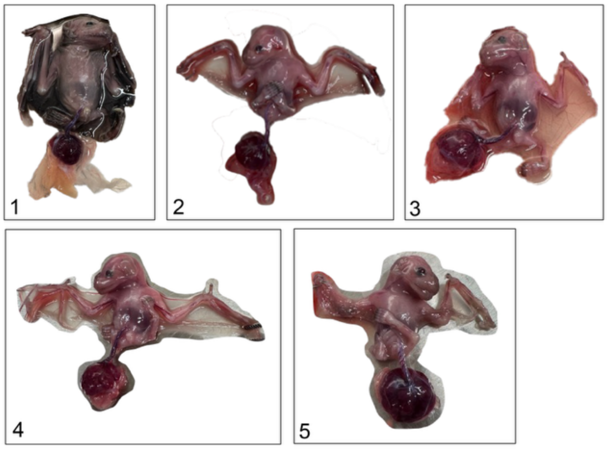


B

| Bat ID # | Placenta Length (cm) | Forelimb Length | CRL (cm) | CRL: Plac ratio | CRL: forelimb ratio | Notable developmental features | Estimated gestational stage |
|---|---|---|---|---|---|---|---|
| 1 | 1.48 | 3.20 | 5.46 | 3.69 | 1.70 | Open eyes, skin pigment and wrinkling, claw pigment, wings opaque, nose-leaf evident. | 24+ (fetal period) |
| 2 | 1.45 | 1.91 | 3.89 | 2.68 | 2.04 | Open eyes, claw pigment, nose-leaf evident, wings translucent. | 21-23 |
| 3 | 1.60 | 1.46 | 4.12 | 2.57 | 2.82 | Open eyes, claw pigment, wings opaque, nose-leaf evident, wings translucent | 21-23 |
| 4 | 1.55 | 1.43 | 3.70 | 2.39 | 2.59 | Open eyes, claw pigment, wings opaque, nose-leaf evident, wings translucent | 21-23 |
| 6 | 1.83 | 1.75 | 3.98 | 2.18 | 2.27 | Open eyes, claw pigment, wings opaque, nose-leaf evident, wings translucent | 21-23 |

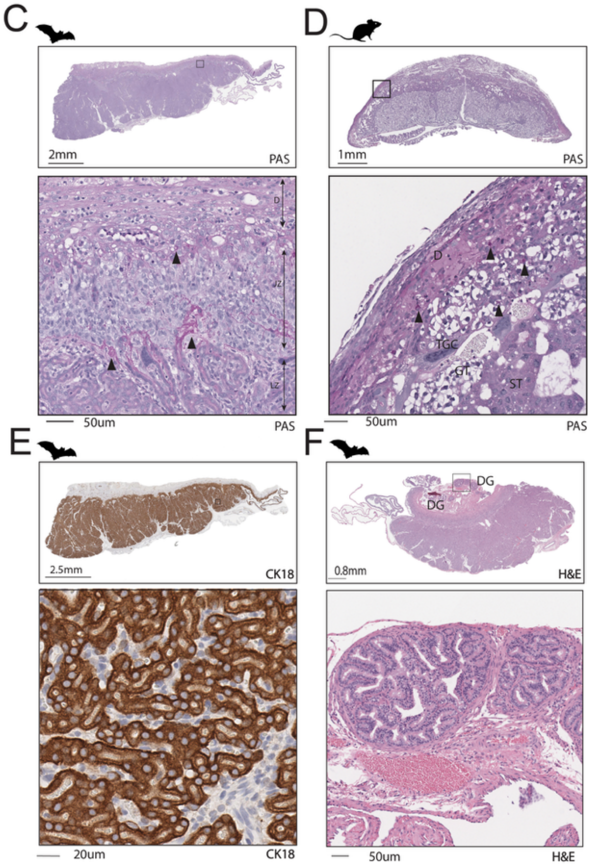


**Figure S1**. **Jamaican Fruit Bat Placental Sampling, Gestational Staging, and Histological Assessment. (A)** Representative photographs of Jamaican fruit bat (Jfb) fetuses at the time of placental isolation, showing a range of developmental stages. **(B)** Table summarizing crown-rump length measurements, morphological features, and estimated gestational ages corresponding to each fetus, based on established Jfb developmental staging criteria. **(C)** Periodic acid–Schiff (PAS) staining of Jamaican fruit bat (Jfb) placenta highlighting glycogen-rich trophoblast regions. **(D)** PAS staining of mouse placenta for comparison, showing typical labyrinth and spongiotrophoblast structures. D (decidua), TGC (trophoblast giant cell), GT (glycogen trophoblast), ST (spongiotrophoblast). **(E)** Immunohistochemistry (IHC) for KRT18 (CK18) in the Jfb placenta section marking trophoblast and epithelial cell populations. **(F)** Hematoxylin and eosin (H&E) staining of the bat decidua showing glandular structures (DG, decidual gland) embedded within the maternal decidua.

**Figure S2**

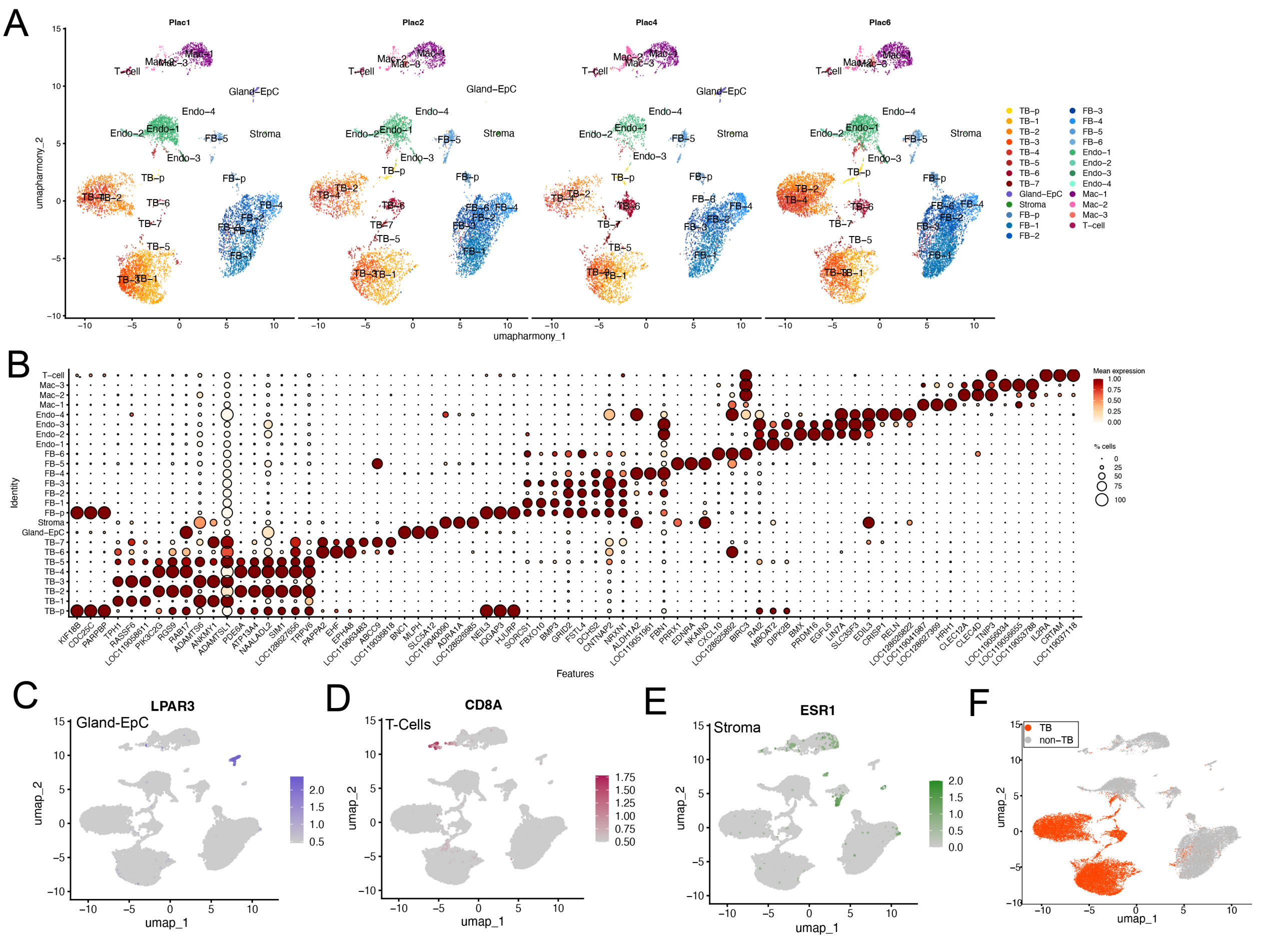


**Figure S2. Defining Cellular Identities of the Jamaican Fruit Bat snRNA-seq dataset. (A)** UMAP split by original sample identity (orig.ident) showing near-equivalent distribution of nuclei across all four biological replicates, confirming lack of major batch effects. **(B)** DotPlot of the top 3 markers per cluster. Scale at right. **(C–E)** Expression of the indicated canonical marker genes across clusters confirming major placental lineages. **(F)** Initial designation of clusters into trophoblast vs. non-trophoblast populations for differential gene expression analysis.

**Figure S3**

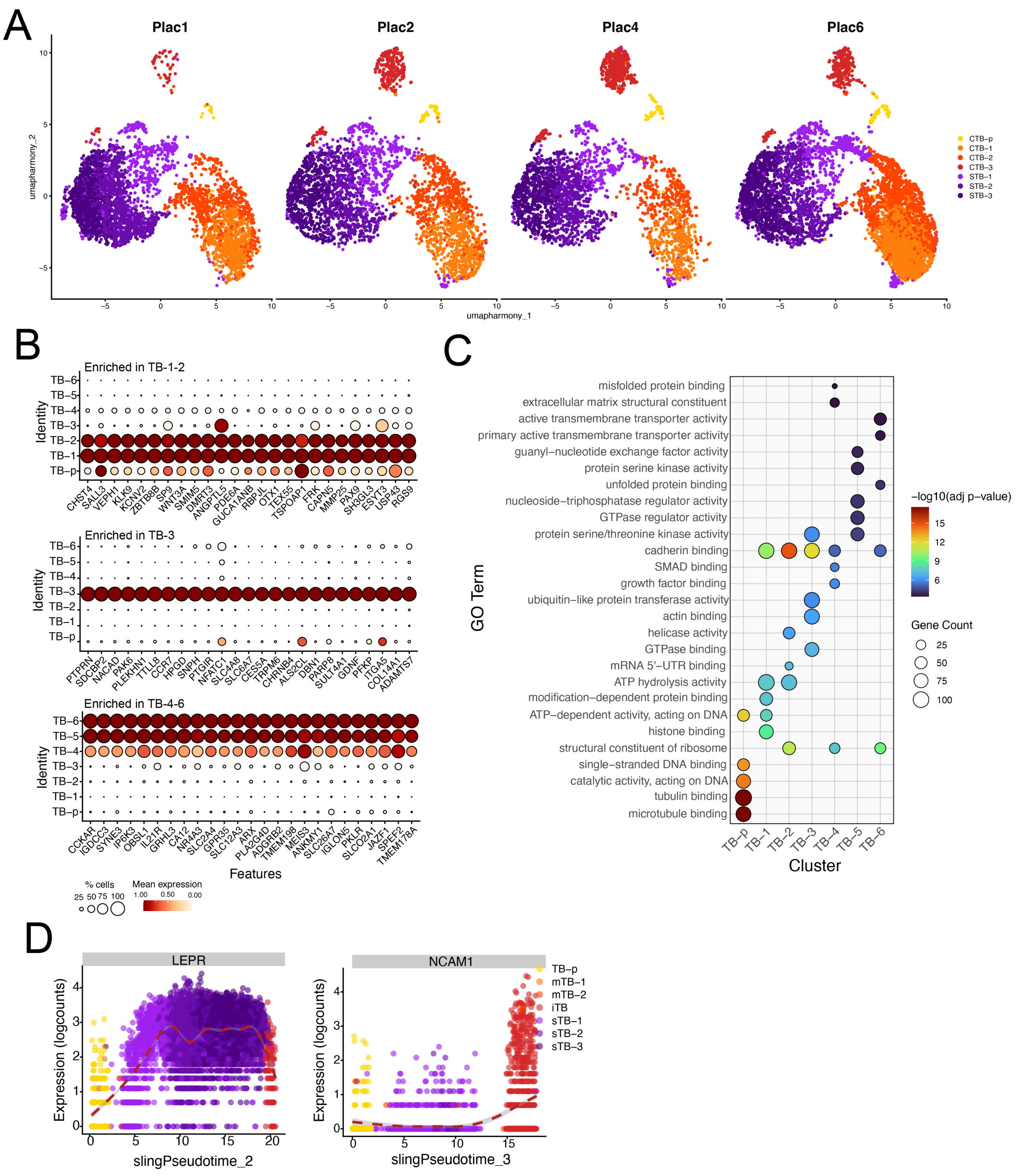

**Figure S3**. **Defining Cellular Identities of Trophoblast Populations in Jamaican Fruit Bat snRNA-seq dataset.** **(A)** Proportion of trophoblast subtypes across biological replicates. **(B)** Heatmap showing cluster-specific expression of key trophoblast-associated genes. **(C)** Gene Ontology enrichment analysis highlighting functional differences between mononuclear, invasive, and syncytial trophoblast populations. **(D)** Gene expression dynamics along inferred Slingshot trajectories for select markers (e.g., LEPR, NCAM1).

#### Figure S4

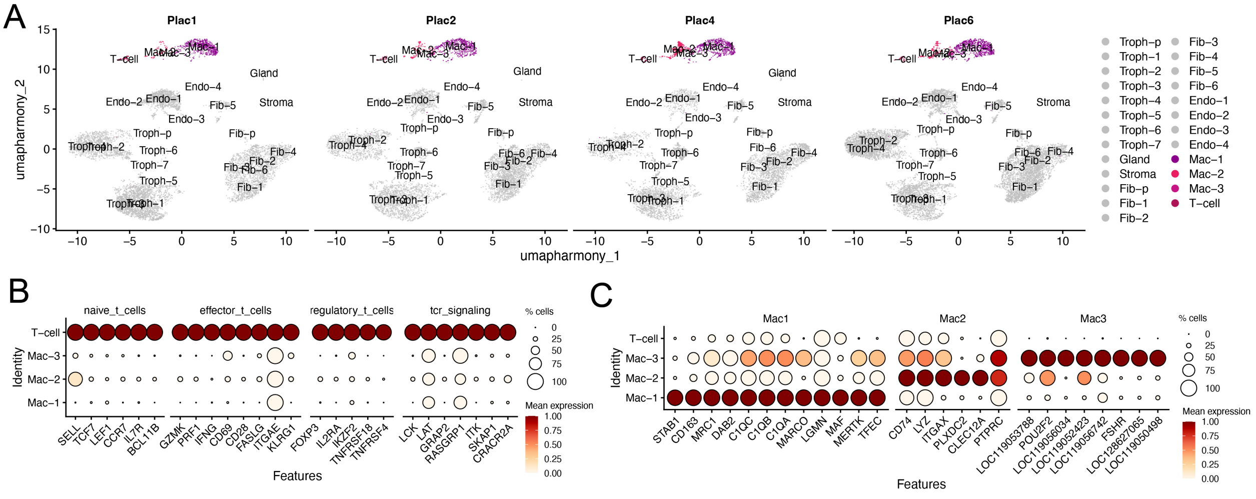


**Figure S4 Defining Cellular Identities of Immune Cell Populations in Jamaican Fruit Bat snRNA-seq dataset. (A)** UMAP split by original sample identity (orig.ident) showing near-equivalent distribution of nuclei across all four biological replicates, confirming lack of major batch effects. Only immune cell subsets are colored. **(B, C)** Expression of canonical markers of T-cells (B) or macrophages (C) across clusters, showing heterogeneity within myeloid and lymphoid compartments.

### Figure S5

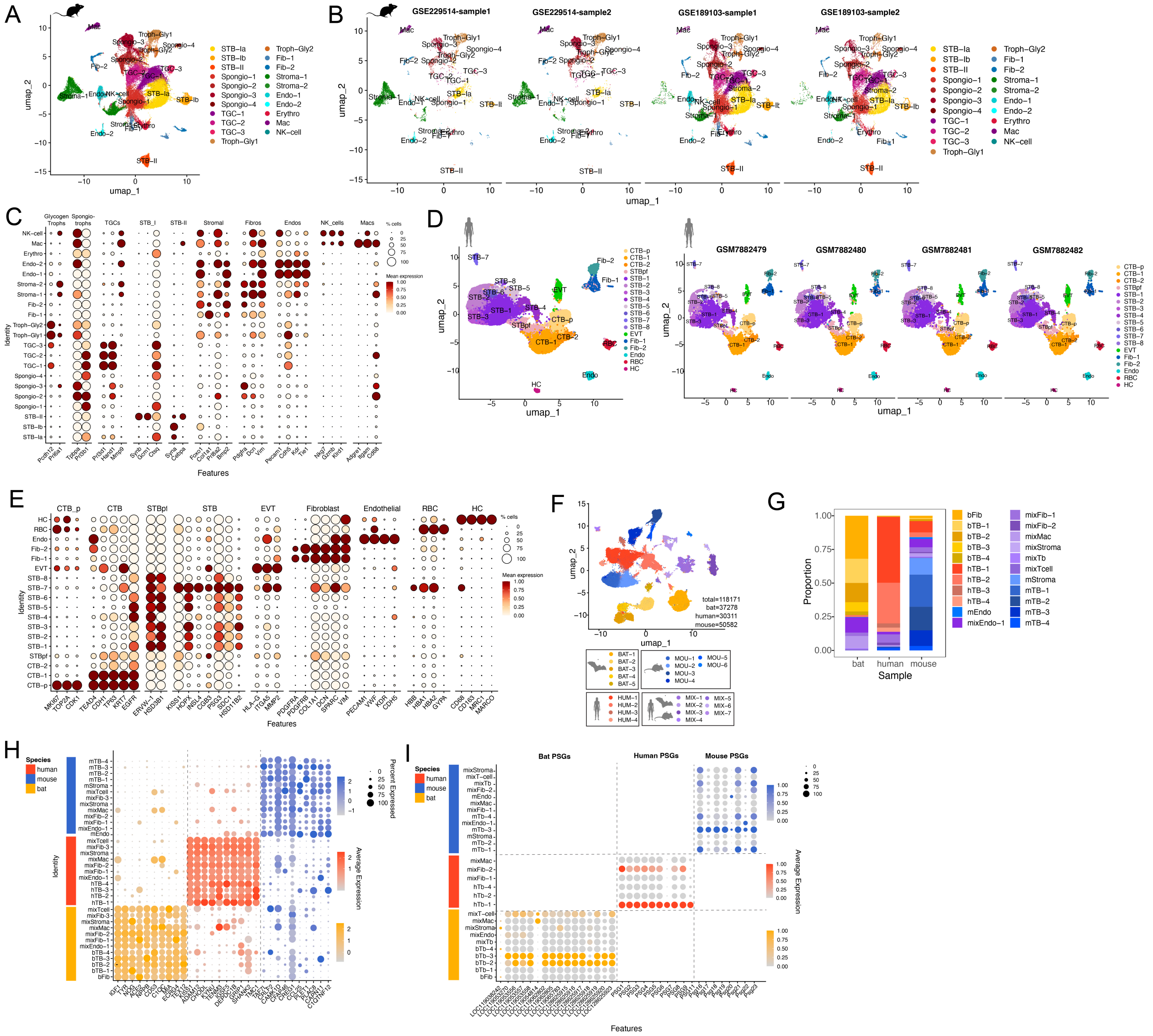


**Figure S5. Cross-Species Integration and Cell Type Validation of Bat, Human, and Mouse Placental Datasets. (A–B)** UMAP visualization of mouse placenta nuclei, shown as a total dataset (A) and split by individual samples (B), validating the presence of major placental cell types based on clustering patterns. **(C)** Dot plot displaying canonical marker expression across identified mouse placental cell populations. **(D)** UMAP visualization of human placenta nuclei, shown as a total dataset (left) and split by individual samples (right), confirming major placental cell populations. **(E)** Dot plot of canonical marker expression across human placental clusters. **(F)** UMAP visualization of 22 integrated clusters across bat, human, and mouse placenta datasets. **(G)** Proportional contribution of each species to individual clusters, highlighting species-specific and mixed-species groups. **(H)** Dot plot showing the expression of the top species-enriched genes in bat (yellow-orange), human (red), and mouse (blue). (I) Dot plot showing the expression of the species-specific PSGs in bat (yellow-orange), human (red), and mouse (blue) clusters.

**Figure S6**

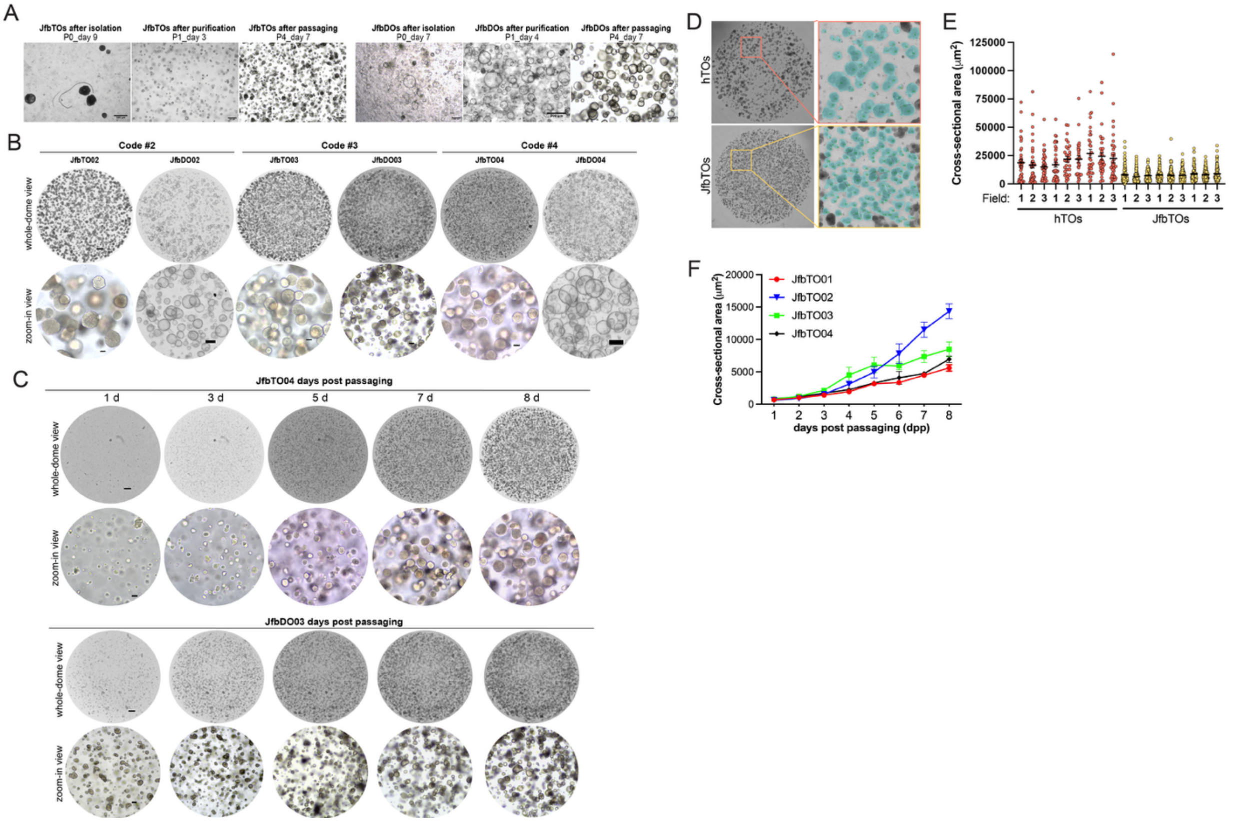


**Figure S6: Imaging and Growth Metrics of Jfb Trophoblast and Decidual gland Organoids. (A)** Representative brightfield images of Jfb TOs (left panels) or DOs (right panels) at the indicated passage number (P) and days post-passaging (day_). Scale bar shown in images. **(B)** Representative brightfield images of three established Jfb TOs or DOs derived from 3 unique placental samples Top row is whole-dome images with scale bar, 500 µm, and bottom are zoomed to show organoid morphology with scale bar, 100 µm for TOs and 200 µm for DOs. **(C)** Representative brightfield images of Jfb TOs (top panels) or DOs (bottom panels) at the indicated days post-passaging. Top row is whole-dome images and bottom are zoomed to show organoid morphology. **(A)** Representative brightfield images in Jfb TOs (left panels) or DOs (right panels) at the indicated passage number (P) and days post-passaging. **(D)** Left panel, representative whole-dome brightfield images of hTOs (top) and JfbTOs (bottom); Right panel, zoomed representative 500 x 500 pixels squares for hTOs (top) and JfbTOs (bottom) cross-sectional area analysis, cyan colored organoid units are the ones chose for analysis based on optimized thresholding. **(E)** Quantification of organoid cross-sectional area across multiple fields of view. Individual datapoints indicate individual organoids. (F) Growth curve of 4 independent JfbTO lines based on organoid cross-sectional area analysis at indicated timepoints.

#### Figure S7

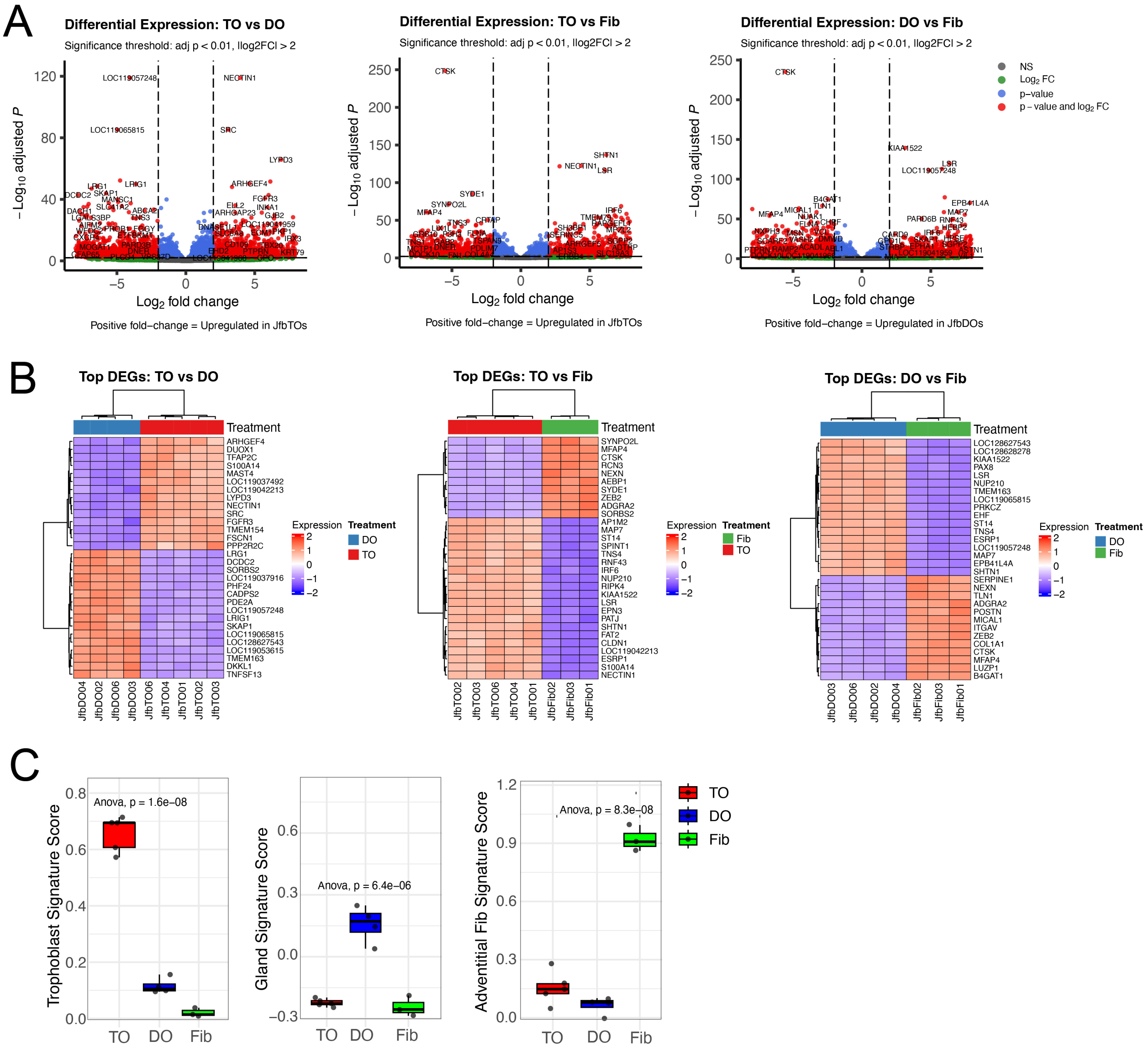


**Figure S7. Bulk RNA-Seq Analysis of Jfb Trophoblast Organoids (TOs), Decidual Organoids (DOs), and Primary Fibroblasts. (A)** Volcano Plots of differentially expressed transcripts in TOs or DOs (left), TOs or fibroblasts (middle), DOs or fibroblasts (right) determined by DeSeq2 analysis. Blue circles represent transcripts that were not significantly changed and red circles represent transcripts significantly enriched, significance was set at $p<0.01$ and $\log_2$fold-change $>\pm 2$. **(B)** Heatmaps (based on $\log_2$ RPKM values) of top differentially expressed transcripts in TOs vs DOs (left), TOs vs fibroblasts (middle), DOs vs fibroblasts (right). Key at right and red indicates high level of expression, purple indicates low levels of expression. Hierarchical clustering is on top. (C) BoxPlots of trophoblast (left), glandular epithelial (middle), or adventitial fibroblasts (right) in TOs (red), DOs (blue), or primary placental fibroblasts (green).

#### Figure S8

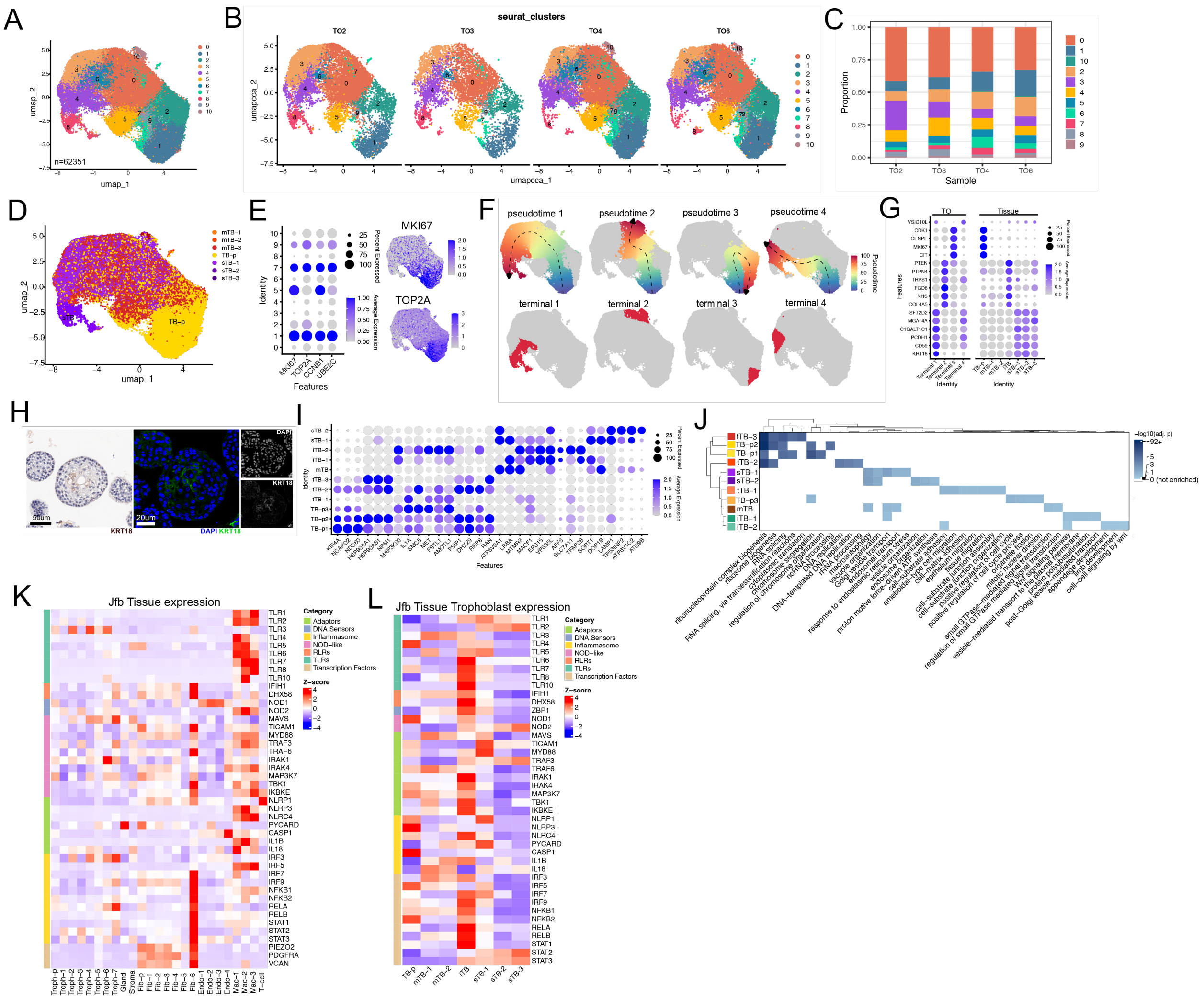


**Figure S8. Defining Cellular Identities of JfbTO snRNA-seq dataset. (A)** UMAP visualization of single-nucleus RNA-seq (snRNA-seq) data from Jamaican fruit bat (Jfb) trophoblast organoids (TOs), identifying 11 transcriptionally distinct clusters. Key at right. **(B–C)** Cluster enrichment across independent TO samples. **(D)** Transfer anchor-based probabilistic projection of placental cell types onto TO snRNA-seq clusters. **(E)** Left, dot plot of proliferation-associated gene expression across TO-derived clusters. Right, feature plots of *MKI67* (top) and *TOP2A* (bottom) highlighting proliferative cells in Cluster 1, designated as the root for trajectory analysis. **(F)** Top, Slingshot trajectory inference revealing four major differentiation paths originating from proliferative progenitors in Cluster 1. Bottom, terminal trajectory states (highlighted in red), representing the terminal 10% of cells along each path. Scale at right. **(G)** Dot plots showing marker gene expression profiles in terminal states, comparing TOs (left) and native placental tissue (right). Key at right **(H)** Left, immunohistochemistry (IHC) for KRT18 in TOs showing localization to internal cavity-lining cells, consistent with syncytialized structures. Right, confocal immunofluorescence staining for KRT18 (green) and DAPI (blue), with grayscale individual channels shown. Scale bar, 20 µm. **(I)** Dot plot of selected differentially expressed genes supporting lineage-specific transcriptional programs. Scale at right. **(J)** Gene Ontology (GO) enrichment analysis across TO-derived trophoblast clusters, revealing functional specialization along the differentiation continuum. Scale at right. **(K, L)** Heatmap showing scaled average expression (Z-score) of key innate immune signaling genes including Toll-like receptors (TLRs), RIG-I–like receptors (RLRs), NOD-like receptors (NLRs), adaptor proteins, inflammasome components, and downstream transcription factors as well as markers of adventitial fibroblasts (in K) across distinct cell populations in the Jfb placenta (K) or tissue-derived trophoblasts (L). Genes are grouped by functional category, with color-coded annotations at left. Expression values were derived from SCT-normalized single-nucleus RNA-seq data and averaged per cluster.

**Supplemental Movie 1**. Overlay of predicted AlphaFold 3.0 structures of a bat pregnancy-specific glycoprotein (PSG6, orange), human PSG1 (red), and mouse Psg17 (blue). The animation highlights the high degree of structural conservation across species, particularly within the immunoglobulin-like domains, despite sequence divergence. Structures are shown aligned to emphasize shared domain architecture and potential functional homology.

**Table S1: Enriched Marker Genes Identified in Jamaican Fruit Bat Placental snRNA-seq.** Marker genes enriched in distinct placental cell populations identified by single-nucleus RNA sequencing of the Jamaican fruit bat placenta. Markers were determined using differential expression analysis with a log2 fold change threshold of log2FC > 0.25 and adjusted p-value adj p < 0.05.

**Table S2: Trophoblast enriched Marker Genes Identified in Jamaican Fruit Bat Placental snRNA-seq.** Marker genes enriched in trophoblast-derived placental cell populations identified by single-nucleus RNA sequencing of the Jamaican fruit bat placenta. Markers were determined using differential expression analysis with a log2 fold change threshold of log2FC > 0.25 and adjusted p-value adj p < 0.05.

**Table S3: Candidate and Re-named Pregnancy-Specific Glycoproteins (PSGs) in the Bat Placenta.** List of bat LOC and CEACAM genes identified as candidate pregnancy-specific glycoproteins (PSGs) based on sequence similarity, transmembrane domain prediction, and structural modeling. Genes confirmed as high-confidence PSGs are re-named PSG1–16 and sequences of these PSGs are shown.

**Table S4: Enriched Marker Genes Identified in subsetted trophoblast populations from Jamaican Fruit Bat Placental snRNA-seq.** Marker genes enriched in distinct trophoblast cell populations that were subsetted and re-clustered from single-nucleus RNA sequencing of the Jamaican fruit bat placenta. Markers were determined using differential expression analysis with a log2 fold change threshold of log2FC > 0.25 and adjusted p-value adj p < 0.05.

**Table S5. Enriched Markers Genes in subsetted Immune Cells from Jamaican Fruit Bat Placental snRNA-seq.** Marker genes enriched in distinct immune cell populations subsetted from single-nucleus RNA sequencing of the Jamaican fruit bat placenta. Markers were determined using differential expression analysis with a log2 fold change threshold of log2FC > 0.25 and adjusted p-value adj p < 0.05.

**Table S6. Jamaican Fruit Bat specific markers in combined Jfb, Human, and Mouse dataset.** Marker genes enriched in Jamaican fruit bat (*Artibeus jamaicensis*) cell populations were identified using the MAST differential expression test applied to the integrated single-nucleus RNA-seq dataset. Genes listed show significant upregulation in bat relative to matched human and mouse clusters, reflecting species-specific transcriptional programs.

**Table S7. Human specific markers in combined Jfb, Human, and Mouse dataset.** Differential expression analysis using MAST identified genes enriched in human placental cell populations relative to corresponding clusters in Jamaican fruit bat and mouse. These markers highlight human-specific transcriptional signatures in trophoblast, stromal, and immune compartments.

**Table S8. Mouse specific markers in combined Jfb, Human, and Mouse dataset.** Genes significantly upregulated in mouse placental cell types compared to human and Jamaican fruit bat were identified using MAST. These mouse-specific markers delineate transcriptional differences that may reflect species-specific placental architecture or lineage specialization.

**Table S9. Bat fibroblast specific markers.** Marker genes enriched in Jamaican fruit bat (Artibeus jamaicensis) fibroblast populations were identified using the MAST test within the integrated cross-species dataset. Genes shown are significantly upregulated in bat-specific fibroblasts relative to shared or homologous fibroblast clusters in human and mouse, highlighting unique aspects of bat stromal gene expression.

**Table S10. DEGs from human TOs treated with polyIC.** DEGs were identified using DESeq2 comparing human TOs treated with polyinosinic:polycytidylic acid (poly I:C) versus mock-treated controls. Genes with significant changes in expression reflect the antiviral and inflammatory transcriptional response of human trophoblasts.

**Table S11. DEGs from Jfb TOs treated with polyIC.** DESeq2 analysis was used to identify genes differentially expressed between poly I:C-treated and mock-treated Jamaican fruit bat TOs. No statistically significant DEGs were detected.

**Table S12. Metadata table of all tissue and organoid experimental materials.** This table summarizes placental tissue samples and trophoblast (JfbTO and hTOs), decidual gland (JfbDO), and Jfb placental primary fibroblast lines used across experiments. For each experiment, the number of distinct tissue samples or cell lines (“codes”) is listed along with the corresponding identifier, estimated gestational stage (in days post-conception, for human as weeks), and organoids/cell tissue culture (TC) passage number. Experiments include single-nucleus RNA-seq (snRNA-seq), bulk RNA-seq, histological staining (H&E), immunohistochemistry (IHC), immunofluorescence (IF), poly I:C treatment, and bright field imaging.
